# Evidence-resolved benchmarks separate biological context recognition from direct-partner specificity in protein interaction models

**DOI:** 10.64898/2026.10.07.757219

**Authors:** Hansen Cheng, Chengyu Sun, Chuan Ye, Chenyang Wu, Dongyu Zhao

**Affiliations:** Department of Biomedical Informatics, School of Basic Medical Sciences, Peking University, Beijing 100191, China; State Key Laboratory of Vascular Homeostasis and Remodeling, Peking University, Beijing 100191, China

## Abstract

Sequence-based models enable scalable protein–protein interaction (PPI) prediction, but whether high scores identify direct physical partners or broader biological compatibility remains unclear. Here we show that biological-context recognition and direct-partner specificity are separable prediction capabilities. We introduce ProtNegDB (http://protnegdb.bjmu.edu.cn/), an evidence-resolved resource distinguishing experimentally supported non-binding from coordinate-verified noncontact within PDB biological assemblies. Using context-matched benchmarks, we find that five sequence-based predictors show limited ability to distinguish direct partners from noncontacting members of the same assembly, whereas PLM-Interact readily ranks both above unrelated proteins. Representation analyses reveal that adaptation to PPI prediction can strengthen biological-context encoding without yielding transferable direct-contact specificity. AlphaFold3 shows greater, though still incomplete, direct-partner resolution, suggesting that explicit multimer modeling can help refine partner specificity. In PLM-Interact, negatives defined by reconstituted direct-binding, cellular binary/proximity and co-complex assays occupy distinct score regimes. Across independent datasets and multiple models, experimental direct non-binding and assembly noncontact further show reproducible, model-dependent scoring patterns. High sequence-based PPI scores should not be interpreted as direct-interface probabilities. Mechanistically interpretable evaluation instead requires explicit consideration of interaction-evidence semantics and partner specificity.

## Introduction

Protein–protein interactions (PPIs) underlie signalling, molecular assembly and cellular regulation, and their systematic prediction remains a major goal of computational biology^1–4^.Sequence-based approaches are particularly attractive because they require only amino acid sequences and support large-scale screening of candidate pairs^5–8^. Recent advances in protein language models (PLMs) have substantially improved sequence-based PPI prediction^9–12^.As these models are increasingly used for candidate prioritization, the biological interpretation of their predictions becomes as important as their benchmark performance. A high interaction score may reflect shared evolutionary, functional or cellular context, but whether it identifies a direct physical partner remains unclear^4,13,14^.

This uncertainty reflects a broader ambiguity in what constitutes a protein interaction. Experimental and database resources aggregate relationships observed at different biological levels, including direct binding between defined components, cellular interaction or proximity, and membership in the same molecular complex^3,4,15–17^.These observations are informative but do not establish the same physical relationship. Sequence representations likewise encode evolutionary and functional information that can make a candidate biologically plausible without specifying whether it forms a direct interface^18–20^. For mechanistic interpretation, complex reconstruction and partner selection, identifying a plausible candidate is insufficient when the question concerns direct physical binding^21–24^. We refer to the ability to identify the direct physical partner of an anchor protein among biologically plausible candidates as direct-partner specificity. Explicit multimer structure prediction provides a complementary approach by modeling candidate inter-chain interfaces and assessing their structural confidence. Biological-context recognition and direct-partner specificity may therefore represent distinct prediction capabilities, although both are commonly evaluated under the broad label of PPI prediction.

Conventional benchmarks make this distinction difficult to assess. Many evaluations contrast known interacting pairs with randomly sampled pairs absent from positive-interaction databases^25–29^.Such controls are better regarded as random unlabeled pairs because database absence does not establish non-interaction, and sampled proteins may differ from true partners in species, sequence cluster, localization or functional context^25,26,30,31^. Such benchmark designs can reward sequence similarity, network topology or other broad compatibility cues and can inflate apparent generalization without establishing whether direct-partner specificity is being measured^27–29,32,33^. Whether these models can resolve the direct physical partner among biologically plausible candidates therefore remains unclear. Experimentally supported negative evidence offers an alternative, but its interpretation depends on the assay, constructs and conditions under which interaction was tested.Purified or reconstituted direct-binding assays test binding between defined components, cellular binary or proximity assays probe relationships in a cellular environment, and co-complex assays detect association that may involve indirect connections^4,15–17,34^. Negative results from these assays therefore constrain different biological hypotheses. Structure-derived noncontact provides another form of evidence in which two proteins coexist within a resolved biological assembly but do not form a direct interface in that structure^30,31^. This observation does not exclude binding in other conformations, assemblies or biological contexts^35–37^. Information leakage from repeated proteins, homologous families and pretrained sequence familiarity presents an additional evaluation challenge^27,28,32,33^. Protein- and cluster-disjoint testing helps assess generalization, but it does not resolve ambiguity in the biological endpoint^27,28,32,33^. Even a fully disjoint benchmark may reward context recognition rather than direct-partner specificity if its negative examples do not distinguish these capabilities. The biological meaning of negative evidence is therefore part of the task definition itself, alongside the separation of training and evaluation data.

Here we introduce ProtNegDB (http://protnegdb.bjmu.edu.cn/), an evidence-resolved resource comprising 11,071 literature-derived negative PPI pairs and 558,112 structure-derived noncontact records from PDB biological assemblies while preserving their distinct evidence semantics. From the structural evidence, we construct 3,365 same-anchor triplets that compare a direct-contact partner with a context-matched noncontacting member of a resolved biological assembly. This design tests direct-partner specificity within a shared biological context. Across five sequence-based predictors, we find limited direct-partner resolution, whereas same-anchor random controls show that PLM-Interact readily ranks both direct partners and assembly noncontacts above unrelated proteins. Representation analyses further show that adaptation to PPI prediction can strengthen biological-context encoding without yielding transferable direct-contact specificity. AlphaFold3 provides greater, although incomplete, direct-partner resolution. In PLM-Interact, negative evidence defined by different experimental configurations occupies distinct score regimes, while experimental direct non-binding and assembly noncontact produce reproducible, model-dependent score differences across independent datasets and multiple models. By separating biological-context recognition from direct-partner specificity, this framework provides an evidence-aware basis for interpreting what PPI model scores represent.

## Results

### ProtNegDB preserves negative-evidence semantics and enables context-matched direct-partner benchmarking

ProtNegDB was constructed from two complementary sources of negative PPI evidence whose biological meanings were retained separately. The literature arm screened a PubTator3 corpus of 40.76 million documents and retained 11,071 literature-derived negative PPI pairs after evidence extraction, staged review and quality control. These included 1,498 pairs supported by direct-binding assays, 761 by cellular interaction assays and 1,836 by co-complex association assays, whereas 6,976 remained assay-unresolved because the available evidence did not support a more specific assignment (Fig. 1a,b). In parallel, the structural arm identified 558,112 coordinate-verified noncontact records between protein chains within PDB biological assemblies. Experimental non-binding and assembly noncontact were retained as distinct evidence types because they constrain different biological relationships. Experimental non-binding is conditional on the assay, constructs and conditions under which interaction was tested, whereas assembly noncontact establishes the absence of a resolved interface only within a particular structural context (Fig. 1a,d).

**Figure 1.**
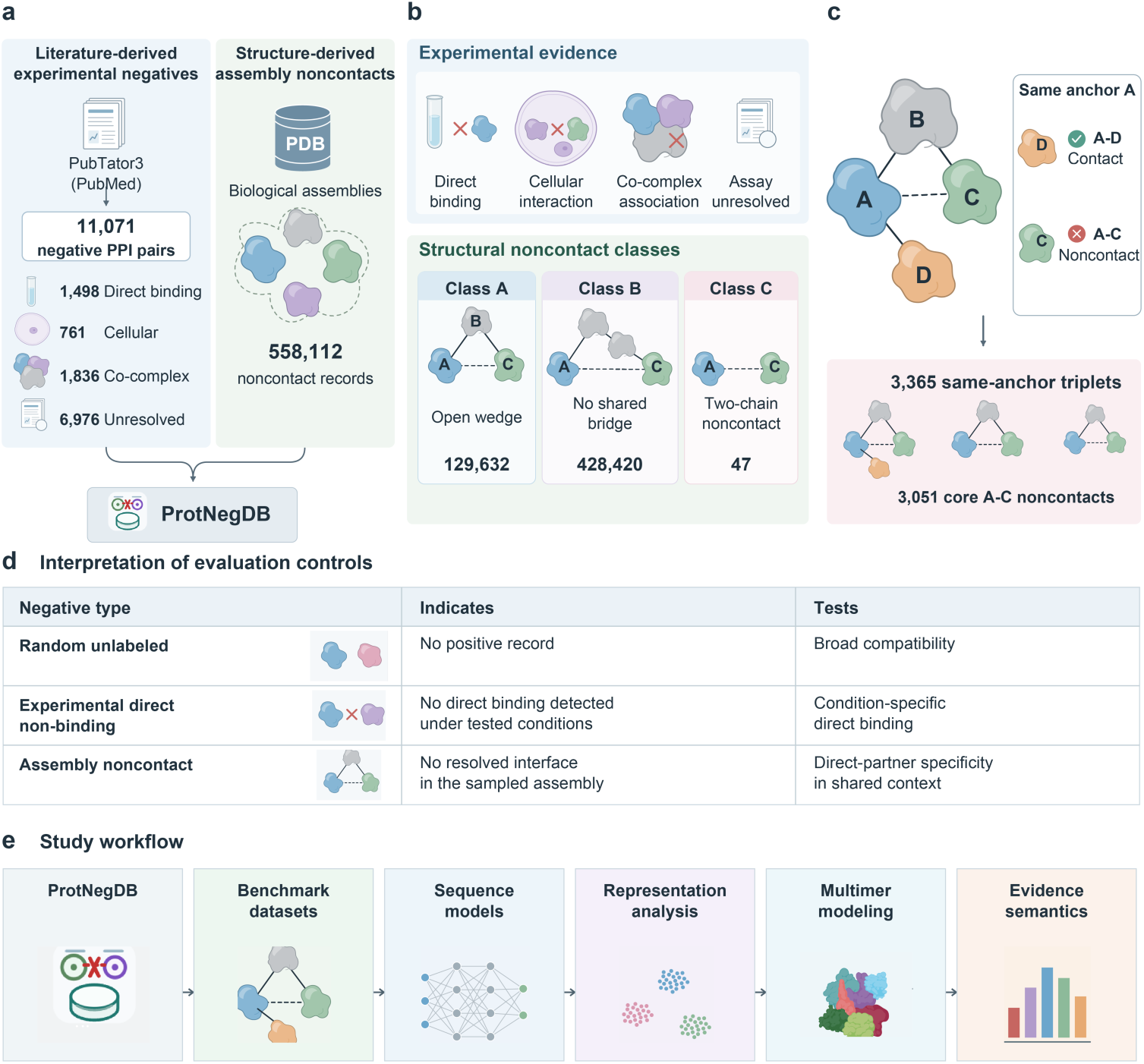
| ProtNegDB preserves negative-evidence semantics and enables context-matched direct-partner benchmarking. **a,** Two complementary sources of negative PPI evidence in ProtNegDB: literature-derived experimental non-binding and structure-derived noncontact within PDB biological assemblies. **b,** Experimental evidence categories and structural noncontact classes. Class A denotes an open wedge in which noncontacting A and C share a contacting bridge B; Class B denotes multichain noncontact without a shared bridge; Class C denotes noncontact in a two-chain assembly. Structural classes describe assembly topology rather than evidence strength. **c,** Same-anchor direct-partner benchmark comparing a direct-contact partner D with an assembly noncontact C for the same anchor A. The primary benchmark contains 3,365 A–D+/A–C− triplets derived from 3,051 core A–C noncontacts. **d,** Biological interpretation of random unlabeled pairs, experimental direct non-binding and assembly noncontact as evaluation controls. **e,** Study workflow linking evidence-resolved negatives to sequence-model evaluation, representation analysis, multimer modeling and negative-evidence semantics.

Structure-derived noncontacts were further classified according to their local topology in the assembly contact graph (Fig. 1b). Class A represents an open wedge in which two noncontacting chains A and C share at least one bridge chain B that contacts both. Class B contains noncontacting pairs from multichain assemblies that contain other contact edges but lack a single shared bridge, whereas Class C comprises noncontacting pairs from assemblies containing exactly two protein chains. These classes describe structural context rather than increasing strengths of biological non-interaction. Applying these definitions yielded 129,632 Class A, 428,420 Class B and 47 Class C records, together with 13 unclassified records.

Class A provides a particularly informative setting for testing direct-partner specificity. In the A–B–C open-wedge configuration, A and C do not form a resolved interface but remain embedded in the same connected molecular environment through a shared contacting chain. C is therefore a context-matched candidate rather than an unrelated negative protein. For each eligible A–C noncontact, we identified a direct-contact partner D of the same anchor A, generating same-anchor A–D+/A–C− comparisons in which the candidates differ in direct-contact status (Fig. 1c). This design asks whether a model can identify the direct physical partner when the competing candidate is already compatible with the anchor’s biological context, rather than merely separating a partner from an unrelated protein (Fig. 1d).

We derived the primary benchmark from the 129,632 resource-level Class A records. A stricter coordinate-resolved branch contained 91,859 candidates before benchmark-specific quality control. Canonical mapping, same-taxon requirements, redundancy control, coordinate filtering and assembly-size stratification yielded 3,051 unique A–C noncontacts from assemblies containing 3–24 protein chains. Among these, 1,606 A–C pairs had at least one eligible D, producing 3,365 same-anchor triplets (Supplementary Fig. S1a,b,e).

The benchmark retained substantial structural support after filtering. Median observed-to-canonical sequence coverage was 0.88 for A and 0.87 for C (Supplementary Fig. S1c). Most A–C pairs were separated substantially beyond the 8 Å contact threshold: 77.7% had a minimum heavy-atom distance of at least 15 Å and 62.7% of at least 20 Å (Supplementary Fig. S1d). Moreover, 1,894 of the 3,051 A–C pairs (62.1%) recurred as noncontacts in at least two distinct PDB entries (Supplementary Fig. S1f). Exact direct-contact conflicts in the PDB pair index were removed during benchmark construction. These observations strengthen the evidence for absence of a direct A–C interface in the sampled structural contexts without implying that the proteins can never interact in another conformation, assembly or condition. ProtNegDB therefore provides complementary experimental and structural negative evidence while preserving the biological endpoint represented by each observation. These evidence-resolved cohorts and same-anchor comparisons form the basis for subsequent analyses of sequence-model prediction, representation geometry, multimer modeling and negative-evidence semantics (Fig. 1e).

### Sequence-based PPI predictors recognize biological context but poorly resolve direct partners

We first asked whether sequence-based PPI predictors could distinguish a direct-contact partner from a biologically plausible but noncontacting candidate. Representative assemblies illustrated the challenge: direct A–D interfaces and noncontacting A–C pairs could receive similarly high PLM-Interact scores despite clear differences in their observed inter-chain geometry (Fig. 2a).

**Figure 2.**
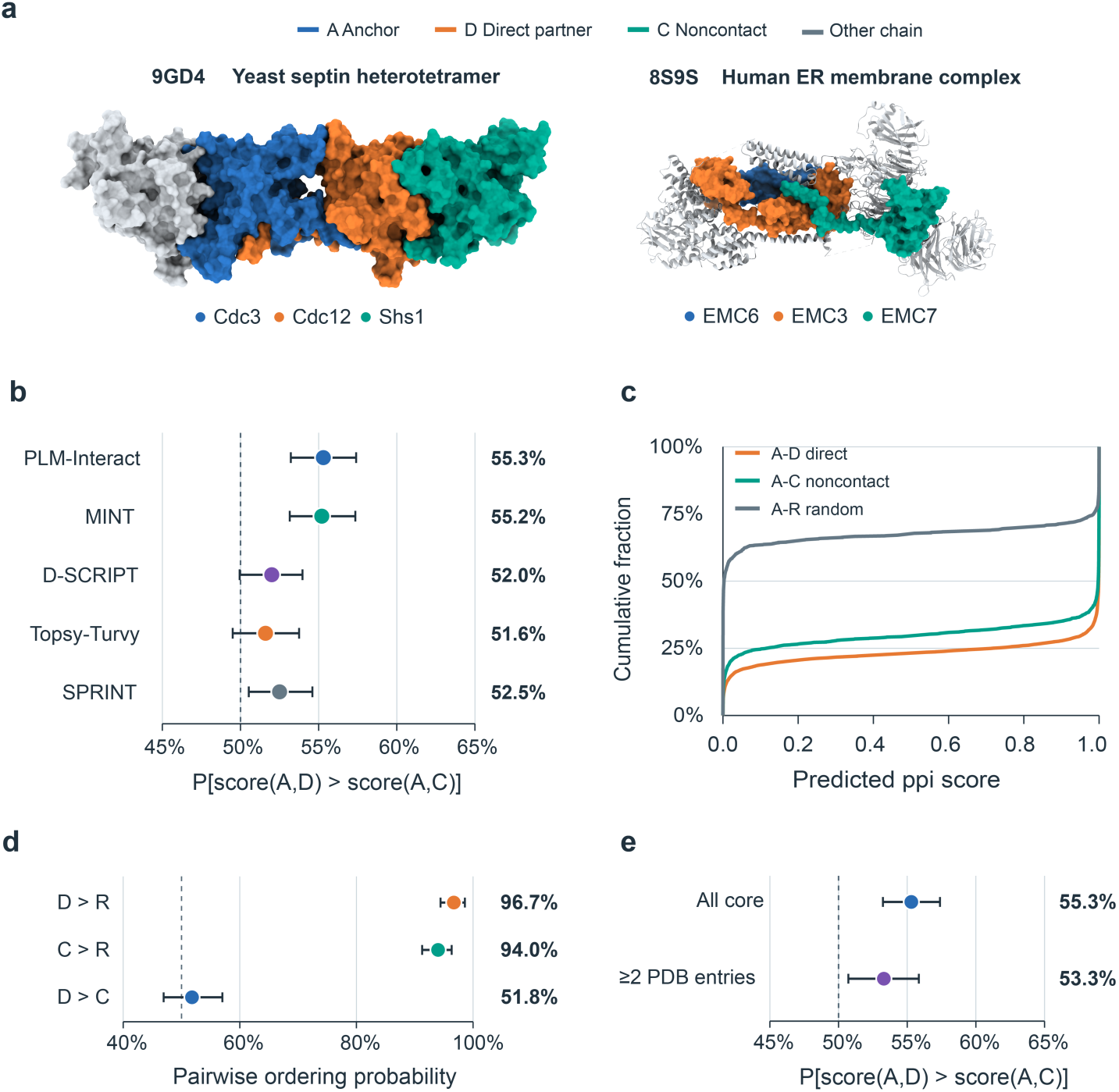
| Sequence-based PPI predictors recognize biological context but poorly resolve direct partners. a, Representative assemblies illustrating direct A–D interfaces and noncontacting A–C pairs assigned similarly high PLM-Interact scores. Other assembly chains are shown in grey. b, Direct-partner ranking across five sequence-based PPI predictors on the complete structural benchmark. The dashed line denotes chance ordering at 0.5. c, PLM-Interact score distributions for direct A–D pairs, assembly-noncontact A–C pairs and random unlabeled A–R pairs. d, Same-anchor random-control analysis. PLM-Interact readily ranks both the direct partner D and assembly noncontact C above an unrelated protein R, while poorly distinguishing D from C. e, Sensitivity of direct-partner ranking to repeated structural support for the A–C noncontact. Error bars denote 95% bootstrap confidence intervals.

We therefore evaluated direct-partner specificity using same-anchor ranking rather than an absolute interaction-score threshold. Across five sequence-based PPI predictors spanning classical and neural sequence-based approaches, the direct partner D outranked the assembly noncontact C in only 51.6–55.3% of comparisons on the complete 3,365-triplet benchmark (Fig. 2b; Supplementary Fig. S2a)^6–10^. The consistently limited ranking across substantially different prediction approaches indicates that the ambiguity is not confined to a single architecture or implementation.

Near-chance D-versus-C ranking did not, however, imply that the models lacked interaction-related information. Direct A–D pairs and Class A assembly noncontacts were both strongly enriched for high PLM-Interact scores, whereas random unlabeled A–R pairs were concentrated at substantially lower scores (Fig. 2c; Supplementary Fig. S2b,c). Thus, PLM-Interact assigned high interaction scores not only to observed direct partners but also to proteins occupying the same biological context without forming the observed interface. To test this distinction directly, we introduced an unrelated same-anchor control. In a locked set of 300 structural triplets representing 266 unique anchors, PLM-Interact ranked D above an unrelated protein R in 96.7% of comparisons and C above R in 94.0%, yet ranked D above C in only 51.8% (Fig. 2d). The model readily recognized both D and C as more compatible with the anchor than an unrelated protein, while providing little information about which compatible candidate formed the observed direct interface. This contrast separates biological-context recognition from direct-partner specificity.

The result was robust to several properties of the structural benchmark. Direct-partner ranking remained close to chance after restricting the analysis to A–C noncontacts observed in at least two distinct PDB entries and across the prespecified 3–10- and 11–24-chain assembly strata (Fig. 2e; Supplementary Fig. S2a,d). Ranking also showed no monotonic dependence on C–D sequence identity (Supplementary Fig. S2e). A conservative provenance analysis further identified 555 triplets in which A–D contact was verified within the same biological assembly supporting the A–C noncontact. Restricting evaluation to this subset did not restore strong direct-partner discrimination across the sequence predictors (Supplementary Fig. S2f). Together, these analyses show that sequence-based PPI predictors can strongly recognize biological compatibility while remaining weak at resolving which plausible candidate forms the direct physical interface.

### PPI adaptation can strengthen biological-context encoding without yielding transferable direct-contact separation

The prediction-level separation between biological-context recognition and direct-partner specificity prompted us to examine whether the same distinction was present in the underlying sequence representations. We analysed frozen representations from ESM2, PLM-Interact and MINT using a locked MMseqs2 50% cluster split (Supplementary Fig. S4)^9,10,20^. Two standardized ridge probes were trained on training-side proteins and evaluated on 540 cluster-unseen triplets. The context probe distinguished biologically plausible A–D and A–C relationships from same-anchor A–R relationships, whereas the contact probe distinguished direct A–D contacts from A–C assembly noncontacts.

Layer-wise analysis revealed a clear divergence between these signals in PLM-Interact. ESM2 representations contained modest transferable context information at the final layer, with a context AUROC of 0.585 (95% triplet-bootstrap CI, 0.568–0.605), whereas direct-contact discrimination remained at chance at 0.500 (0.485–0.513). After adaptation for PPI prediction, PLM-Interact increased context separation to 0.708 (0.681–0.731), but contact separation remained essentially unchanged at 0.507 (0.480–0.530) (Fig. 3a; Supplementary Fig. S3a). The strongest representational change was therefore increased separation of biologically plausible A–D/A–C relationships from unrelated A–R pairs rather than separation of A–D from A–C.

**Figure 3.**
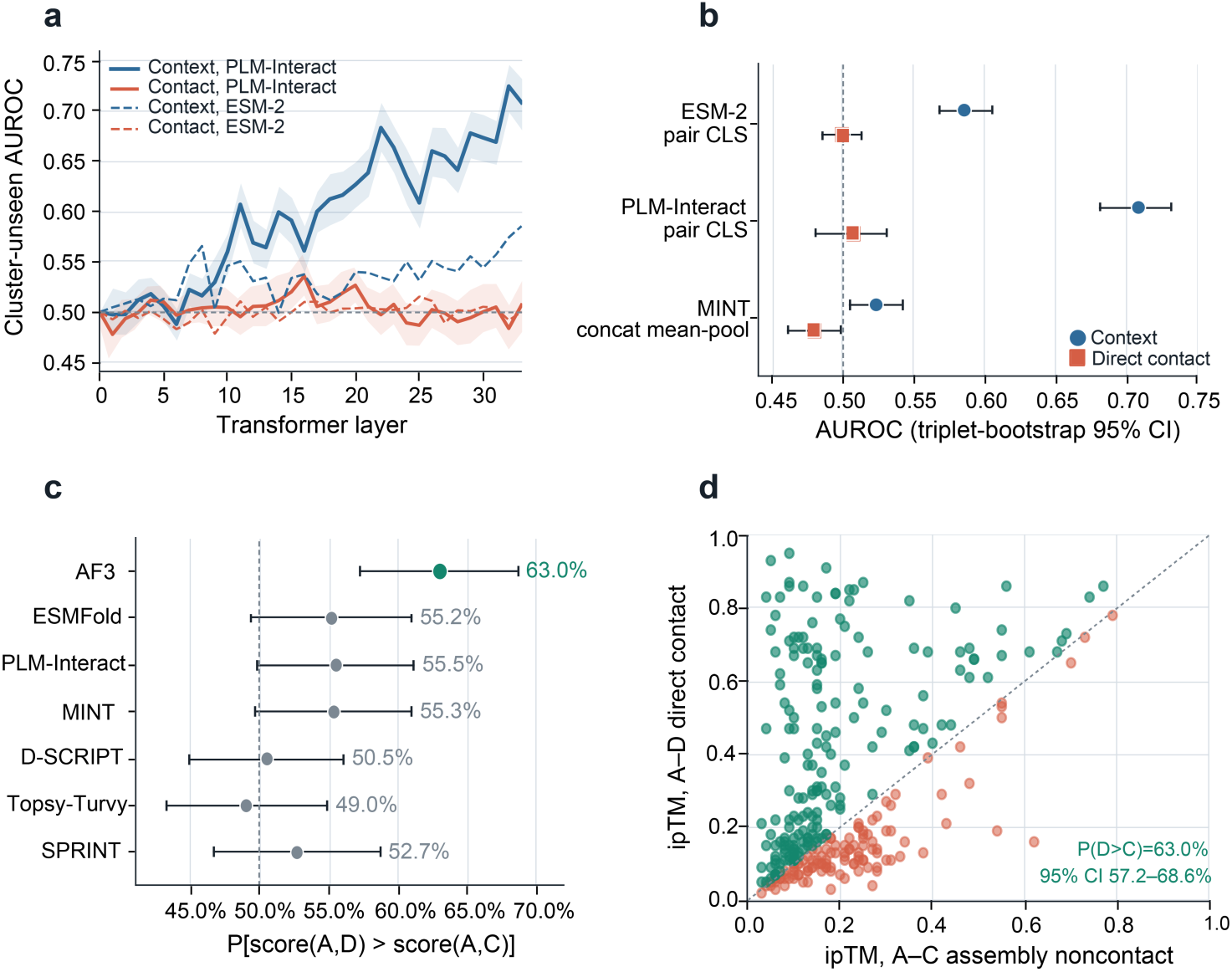
| PPI adaptation preserves context ambiguity, whereas AlphaFold 3 provides partial direct-partner resolution. **a,** Cluster-unseen context and direct-contact probe performance across ESM2 and PLM-Interact layers. The context probe distinguishes biologically plausible A–D/A–C pairs from A–R pairs; the contact probe distinguishes A–D from A–C. **b,** Final-layer context and contact probe performance for ESM2, PLM-Interact and MINT. Error bars denote triplet-bootstrap 95% confidence intervals. **c,** Direct-partner ranking of AlphaFold 3, ESMFold and five sequence-based predictors on the common 300-triplet, 256-anchor subset. The dashed line denotes chance ordering. **d,** Paired AlphaFold 3 ipTM scores for the direct A–D and assembly-noncontact A–C candidate complexes. Points above the diagonal indicate higher ipTM for the direct partner. Confidence intervals in **c,d** use anchor-block bootstrap resampling.

The final-layer comparison across representation architectures showed the same qualitative asymmetry (Fig. 3b). MINT achieved a context AUROC of 0.523 (0.505–0.541) and a contact AUROC of 0.479 (0.461–0.498). Layer-wise and subspace analyses likewise failed to reveal strong transferable direct-contact separation (Supplementary Fig. S3b,c,e). Consistent with these results, same-anchor representation geometry in PLM-Interact placed A–D and A–C substantially closer to one another than A–D and A–R, with final-layer cosine similarities of 0.784 and 0.368, respectively (Supplementary Fig. S3d).

To test whether explicit structural-noncontact supervision could make this distinction transferable, we fine-tuned PLM-Interact on matched datasets containing identical anchors, positive partners and negative-protein multisets. Structure-FT retained the observed A–C pairings, whereas Random-FT reassigned the same negative proteins within taxon. Although the structural pairings were learnable in the exposed training distribution, no detectable advantage transferred to the 540 cluster-unseen triplets. Direct-partner ranking was 0.521 for the original model, 0.523 after Random-FT and 0.526 after Structure-FT, with the same conclusion across seeds and stringent sensitivity analyses (Supplementary Fig. S5a–f). Under the present training regime and dataset size, structural-negative supervision alone therefore did not yield a measurable cluster -unseen advantage over matched random-negative training. At the representation level, PPI adaptation was asymmetric, strengthening transferable context encoding without a corresponding gain in direct-contact separation.

### AlphaFold 3 adds partial direct-partner resolution beyond sequence-level models

We then tested whether explicit candidate-complex modeling could better distinguish a direct partner from a context-matched assembly noncontact. AlphaFold 3, ESMFold and the five sequence-based predictors were compared on a locked common subset of 300 triplets representing 256 unique anchors^20,21^. AlphaFold 3 showed the highest direct-partner ranking point estimate, placing D above C in 63.0% of comparisons. ESMFold reached 55.2%, while the five sequence-based predictors ranged from 49.0% to 55.5% (Fig. 3c). Thus, AlphaFold 3 provided greater partner resolution at the point-estimate level, although the separation remained far from complete.

The paired AlphaFold 3 analysis showed both the improvement and its limitation. AlphaFold 3 assigned a higher ipTM to the A–D candidate complex than to A–C in 63.0% of triplets (95% CI, 57.2–68.6%; Fig. 3d). The distributions remained substantially overlapping, and many A–C noncontacts received ipTM values comparable to or greater than those of the corresponding direct partners (Supplementary Fig. S6a,b).

The common-set comparison should therefore be interpreted as greater but incomplete direct-partner resolution rather than universal statistical superiority over every sequence predictor. Paired differences were clearly resolved against several baselines, whereas uncertainty remained for some of the stronger sequence-based comparators (Supplementary Fig. S6c). Moreover, this benchmark does not establish formal zero-shot generalization because exact historical pair exposure could not be reconstructed.

Explicit candidate-complex modeling adds partner-specific information beyond the sequence-level predictions examined here, but substantial direct-partner ambiguity remains.

### Different negative-evidence semantics occupy distinct score regimes in PLM-Interact

The strong scores assigned to assembly noncontacts raised a broader question: whether negative pairs defined by different experimental and structural evidence represent the same prediction endpoint. We harmonized Negatome records according to the biological relationship tested by the underlying assay, yielding 298 reconstituted direct non-binding pairs, 145 cellular binary or proximity negatives and 119 co-complex association negatives after common quality control (Supplementary Fig. S7a,c)^30,31^. These cohorts were compared with 2,480 clean assembly noncontacts and 2,000 random unlabeled pairs.

The resulting PLM-Interact predictions differed markedly across evidence classes (Fig. 4a). Using 0.5 as the model’s binary operating threshold, only 16.1% of reconstituted direct non-binding pairs were classified as interaction-positive, compared with 42.1% of cellular binary or proximity negatives and 52.9% of co-complex association negatives^9^. Assembly noncontacts showed the highest positive-call fraction at 71.4%, whereas 32.5% of random unlabeled pairs exceeded the same threshold. Full score distributions and filtering sensitivities supported the same qualitative separation (Supplementary Fig. S7b–d).

**Figure 4.**
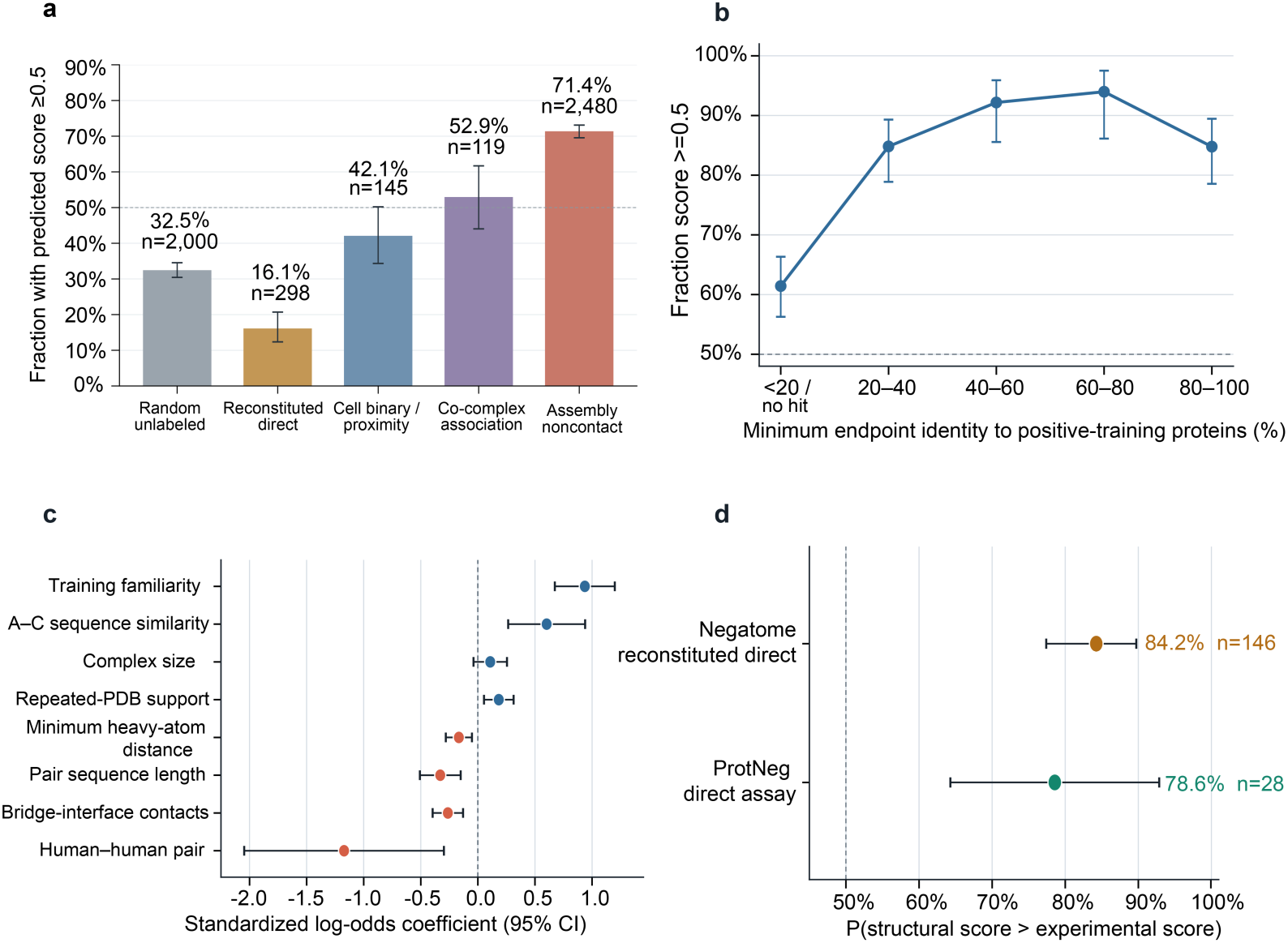
| Negative-evidence semantics correspond to distinct PLM-Interact score regimes beyond training familiarity. **a,** Fraction of pairs receiving an interaction-positive PLM-Interact score (≥0.5) across random unlabeled pairs, reconstituted direct non-binding, cellular binary/proximity negatives, co-complex negatives and assembly noncontacts. For evidence-defined negative sets, lower fractions indicate fewer apparent positive calls for the corresponding endpoint; the categories do not represent a common false-positive rate because they test different biological relationships. **b,** Fraction of assembly noncontacts scoring ≥0.5 stratified by minimum endpoint similarity to proteins observed in positive training interactions. **c,** Multivariable associations with interaction-positive scoring among assembly noncontacts. Points show standardized log-odds coefficients and 95% confidence intervals. **d,** Probability that an assembly noncontact receives a higher PLM-Interact score than a familiarity- and length-matched experimental direct-negative pair. Results are shown independently for Negatome and ProtNegDB direct negatives.

These proportions should not be interpreted as false-positive rates for a single universal interaction task. Reconstituted direct-binding assays test binding between defined components under specified conditions; cellular binary or proximity assays probe relationships in a cellular environment; co-complex measurements can capture indirect association; and assembly noncontact denotes absence of an observed interface within a particular resolved structural context. A high score for an assembly noncontact is therefore an apparent false positive only if the score is interpreted specifically as a probability of direct physical contact. The ordering in Fig. 4a reflects differences in evidence semantics rather than a hierarchy of negative-label quality.

Because assembly noncontacts were especially enriched for high scores, we next asked whether this pattern could be explained by sequence familiarity with proteins represented in positive training interactions. The fraction scoring at least 0.5 increased strongly with endpoint similarity to positive-training proteins (Fig. 4b). Training familiarity remained the strongest positive correlate in a multivariable analysis that additionally considered A–C sequence similarity, complex size, repeated-PDB support, minimum heavy-atom distance, pair length, bridge-interface contacts and species composition (Fig. 4c). Alternative familiarity definitions, overlap handling and uncertainty estimators supported the same conclusion (Supplementary Fig. S8a–e).

Training familiarity nevertheless did not account for the experimental–structural separation. After matching human assembly noncontacts to experimental direct negatives on endpoint familiarity and sequence-length characteristics, the structural pair received the higher PLM-Interact score in 84.2% of 146 comparisons against Negatome direct negatives and 78.6% of 28 comparisons against reviewed ProtNegDB direct-assay negatives (Fig. 4d).

Training familiarity modulates PLM-Interact scores but does not account for the broader separation among evidence-defined negative classes, reinforcing the need to treat negative-evidence semantics as part of the prediction endpoint.

### Experimental–structural score differences are reproducible across datasets but model-dependent

To test whether the experimental–structural separation generalized beyond historical Negatome records, we examined an independently reviewed ProtNegDB cohort comprising 480 accepted negative PPI pairs from 362 publications. After taxon-aware mapping, sequence retrieval and removal of exact Negatome overlaps, 104 pairs supported by biophysical or other direct-binding assays formed the primary contemporary direct non-binding cohort (Supplementary Fig. S9a,b).

In PLM-Interact, these contemporary direct negatives reproduced the Negatome pattern. Only 25.0% scored at least 0.5, with a median score of 0.0079, compared with 16.1% and a median of 0.0003 for Negatome reconstituted-direct negatives and 71.4% and a median of 0.9976 for assembly noncontacts (Fig. 5a). The full score distributions showed the same separation, indicating that the result did not depend on a particular binary threshold.

**Figure 5.**
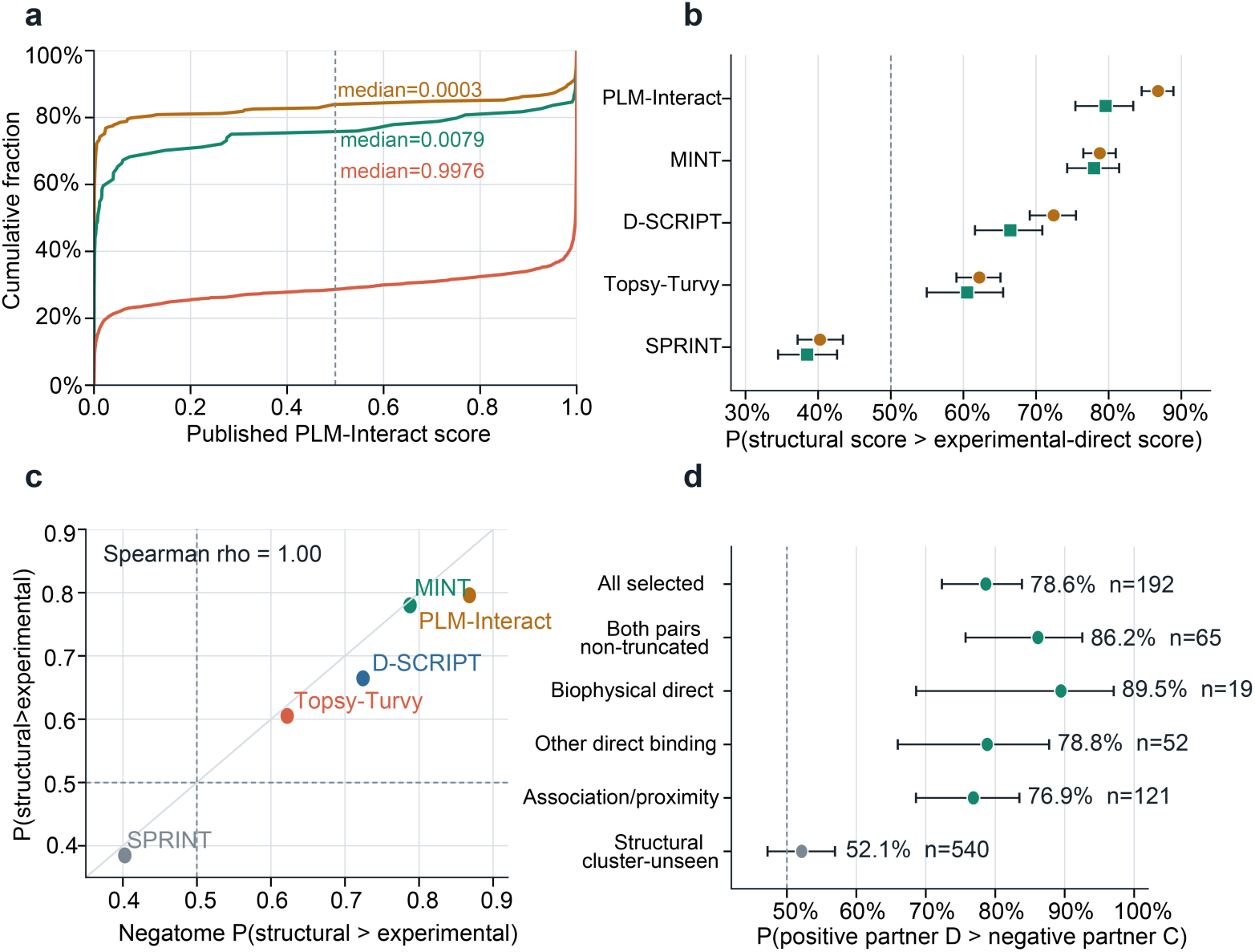
| Experimental–structural score differences reproduce across datasets but vary across PPI models. **a,** PLM-Interact score distributions for independently curated Negatome and ProtNegDB direct non-binding pairs compared with assembly noncontacts. **b,** Probability that an assembly-noncontact score exceeds an experimental-direct score across five PPI predictors. Negatome and ProtNegDB provide independent experimental-negative cohorts; 0.5 denotes no systematic ordering. **c,** Cross-dataset concordance of the model-specific experimental–structural ordering in Negatome and ProtNegDB. Each point represents one predictor; dashed lines denote 0.5 and the diagonal denotes equal effects across datasets. **d,** Supportive PLM-Interact ranking of IntAct-annotated direct-positive partners against reviewed ProtNegDB negative partners, including assay and sequence-completeness strata. The structural cluster-unseen benchmark is shown for reference. Error bars denote 95% confidence intervals.

We next compared experimental-direct and structural-noncontact scores across five sequence-based predictors using the probability that a structural score exceeded an experimental-direct score. With Negatome negatives, this probability was 0.868 for PLM-Interact, 0.788 for MINT, 0.724 for D-SCRIPT and 0.622 for Topsy-Turvy. The independent ProtNegDB cohort reproduced the same orientation, with corresponding values of 0.796, 0.780, 0.665 and 0.605 (Fig. 5b). SPRINT showed the opposite pattern in both datasets, with probabilities of 0.403 and 0.385, respectively. Experimental–structural score separation was therefore reproducible within individual models, but its magnitude and even direction differed across models.

This model-specific behaviour was highly concordant between the two independently curated datasets. The rank ordering of the experimental–structural effect across the five predictors was identical between Negatome and ProtNegDB (Spearman ρ = 1.00; Fig. 5c). Importantly, the reversed ordering in SPRINT did not translate into improved direct-partner specificity: SPRINT remained near chance when ranking D against a context-matched C in the structural benchmark (Fig. 2b). Population-level ordering of negative-evidence classes and same-anchor direct-partner discrimination therefore represent distinct properties of a PPI model.

As a complementary analysis, PLM-Interact ranked an IntAct MI:0407-annotated direct-positive partner above the reviewed negative partner in 78.6% of 192 eligible comparisons, increasing to 86.2% when neither pair required sequence truncation and to 89.5% for biophysical-direct negatives (Fig. 5d)^17^. Sensitivity analyses across truncation, species and positive-database support showed the same overall direction (Supplementary Fig. S9c–e). Because the IntAct positive candidates were not individually re-curated from their primary publications, this analysis is supportive rather than a primary independent validation.

Reproducibility across datasets, coupled with model-specific magnitude and direction, argues against a universal numerical interpretation of PPI scores. Their meaning depends on both the evidence endpoint and the model that produces them.

## Discussion

Our results identify two sources of ambiguity in protein–protein interaction prediction. First, biological-context recognition and direct-partner specificity are separable capabilities. Across five sequence-based predictors, context-matched assembly noncontacts were difficult to distinguish from direct-contact partners, even though PLM-Interact readily ranked both above unrelated proteins. Second, negative evidence is not a single biological class: direct-binding, cellular interaction, co-complex association and assembly noncontact represent different endpoints and occupy distinct model-score regimes. These two forms of ambiguity caution against interpreting a high PPI score as a universal probability of direct physical binding.

The same-anchor benchmark makes this distinction explicit. Random negatives can differ from true partners in many properties beyond direct contact and may therefore reward broad biological compatibility^25–29^. By contrast, assembly noncontacts preserve much of the molecular context while differing in observed interface status. PLM-Interact ranked D above unrelated proteins in 96.7% of comparisons and C above unrelated proteins in 94.0%, yet ranked D above C in only 51.8%. Thus, the model clearly recognizes biologically plausible partners but provides limited information about which candidate forms the direct interface. Direct-partner specificity should be evaluated separately when predictions are used for mechanistic interpretation, partner selection or complex reconstruction.

Representation analyses support this interpretation. PPI adaptation from ESM2 to PLM-Interact increased transferable context separation without producing comparable direct-contact separation, suggesting that PPI-oriented training can emphasize biological compatibility more readily than interface identity. This conclusion should not be taken to imply that sequence representations contain no contact-related information; nonlinear, residue-level or cluster-specific signals may not be accessible to the probes used here. Likewise, the lack of cluster-unseen improvement after structural-negative fine-tuning does not show that such supervision is intrinsically ineffective, but indicates that under the present training regime it was insufficient to produce transferable partner specificity. AlphaFold 3 provided greater direct-partner resolution than the sequence-level models at the point-estimate level, consistent with explicit candidate-complex modeling contributing additional partner-specific information^21–24^. However, its 63.0% ranking accuracy and substantial overlap between direct and noncontacting candidates show that this ambiguity is reduced rather than eliminated.

A second implication concerns the meaning of negative interaction evidence. Reconstituted direct-binding assays, cellular binary or proximity assays, co-complex measurements and structural noncontact do not test equivalent hypotheses^15,30,31^. Their different PLM-Interact score distributions should therefore not be interpreted as a hierarchy of negative quality. For a reconstituted direct non-binding pair, an interaction-positive prediction can reasonably be viewed as an apparent false positive for direct binding. For an assembly noncontact, the same score may instead reflect broader biological compatibility despite the absence of an interface in the resolved structure. Training familiarity increased structural-noncontact scores, but matching and multivariable analyses showed that it did not explain the experimental–structural separation. The biological semantics of the negative evidence are therefore part of the prediction task itself.

This evidence-semantic effect was reproducible across independently curated datasets but was not universal across models. PLM-Interact, MINT, D-SCRIPT and Topsy-Turvy consistently assigned higher scores to assembly noncontacts than to experimental direct negatives, whereas SPRINT reproducibly showed the opposite orientation. Importantly, SPRINT remained near chance in the context-matched partner-selection benchmark, showing that population-level ordering of negative-evidence classes is distinct from direct-partner specificity. PPI scores therefore do not lie on a model-independent biological scale: their interpretation depends both on the model and on the biological endpoint represented by the evaluation data.

Several limitations define the scope of these conclusions. Assembly noncontact is structure-and condition-specific and cannot establish that two proteins never interact in another conformation, stoichiometry or biological state^35–37^. Experimental negatives are likewise conditional on assay design and constructs. The representation analyses assess transferable linear accessibility rather than all information potentially encoded by a model, and the AlphaFold 3 comparison used a smaller common benchmark and an ipTM score that should not be treated as a calibrated binding probability. Nevertheless, the results converge on a common principle: PPI prediction comprises multiple related but non-equivalent tasks. Future benchmarks should distinguish broad biological compatibility from direct-partner specificity and preserve the biological meaning of negative evidence rather than collapsing all absent interactions into one class. Doing so should make PPI model evaluation more mechanistically interpretable and better aligned with the biological questions for which these models are used.

## Methods

### Study design and evidence semantics

We treated negative PPI evidence according to the biological observation that generated it and did not collapse literature-derived experimental non-binding, structure-derived assembly noncontact and random unlabeled pairing into a single negative class. Experimental direct non-binding denotes failure to detect binding under the constructs, assay and conditions reported in the source publication. Assembly noncontact denotes absence of a resolved direct interface between two protein chains in a specified PDB biological assembly. Random A–R controls were used only as unlabeled reference pairs and were not interpreted as demonstrated biological non-interactions.

The primary structural task was same-anchor direct-partner discrimination. Each triplet shared an anchor A and contrasted a direct-contact partner D with a candidate C that did not contact A in a specified biological assembly. The biological-context constraint therefore applies directly to C, whereas D was supported by an observed A–D contact either in the same assembly or in the aggregate PDB contact index. A conservative sensitivity analysis separately restricted evaluation to triplets with verified same-assembly A–D contact. Unless otherwise stated, exact ranking ties contributed 0.5.

### ProtNegDB literature-evidence construction

The literature arm was constructed from a local PubTator3 BioCXML corpus comprising 40,763,455 document records distributed across ten BioCXML shards^38^. PubMed and PMC document categories overlap and were therefore not treated as additive source counts. Entity-aware prescreening identified 29,418,388 records containing annotated entities. Documents containing at least two distinct gene entities produced 6,021,850 candidate PPI articles; a broader gene/chemical candidate universe contained 8,699,676 records.

Negative-interaction trigger mining identified approximately 948,877 candidate sentences. Expansion across sentence-level entity pairs produced 2,805,068 sentence–entity-pair evidence rows from 529,977 articles. All candidate articles entered an article-level LLM-assisted evidence review, generating 354,311 initial claims. Aggregation across redundant evidence produced 345,406 claims representing 266,711 unique candidate pairs.

A subsequent pair-level review considered 277,316 candidate pairs and classified each as retain, human-review or discard according to the reported experimental evidence and claim scope. The retained set contained 43,843 pairs, including 19,087 protein–protein and 24,692 protein–small-molecule candidates. A final stringent evidence review retained records consistent with physical-negative evidence or with an explicitly documented scope restriction. The main ProtNegDB literature PPI resource contained 11,071 negative PPI pairs.

Experimental methods were classified according to assay configuration rather than keyword alone. Direct-binding assays included biophysical measurements such as SPR, ITC, BLI, MST and fluorescence anisotropy, together with appropriately configured purified pull-down, in-vitro binding and far-western assays. Cellular interaction assays included Y2H, BRET, FRET, BiFC, PLA and related cellular binary/proximity measurements. Co-complex association included co-immunoprecipitation and affinity-based complex-detection assays. Records for which the available evidence did not support assignment to these categories were retained as assay-unresolved. The final resource contained 1,498 direct-binding, 761 cellular-interaction, 1,836 co-complex-association and 6,976 assay-unresolved pairs.

### ProtNegDB structural-evidence construction

Structural evidence was derived from RCSB PDB biological assemblies rather than asymmetric units^39^. Candidate entries were retrieved as pre-expanded assemblyN coordinate files. The source query contained 197,375 experimental biological assemblies with at least two protein chains.

Protein-chain contact was defined using minimum heavy-atom distance. Two chains were classified as contacting if any inter-chain heavy-atom distance was ≤8 Å and as noncontacting if all inter-chain heavy-atom distances exceeded 8 Å. Contact graphs were constructed separately for each biological assembly.

Assembly noncontacts were classified by local contact topology. Class A contained open wedges in assemblies with at least three protein chains, such that A and C did not contact one another but at least one bridge chain B contacted both. Class B contained noncontacting pairs in multichain assemblies that contained other contact edges but lacked a single bridge contacting both A and C. Class C contained noncontacting pairs from assemblies containing exactly two protein chains. Records that passed the resource-level noncontact criteria but lacked a supported topological assignment were retained as unclassified.

The final structural resource comprised 558,112 assembly-noncontact records: 129,632 Class A, 428,420 Class B, 47 Class C and 13 unclassified records. These are assembly-level structural evidence records and should not be interpreted as unique universal non-interacting UniProt pairs.

### Class A benchmark construction

For direct-partner benchmarking, we used a stricter coordinate-resolved branch of 91,859 Class A open-wedge records. Chains were mapped to canonical UniProt accessions and sequences^40^. Candidate A and C proteins were required to originate from the same reported taxon and to map to distinct protein chains. Both endpoints required at least 30 observed residues, observed-to-entity construct coverage ≥0.5 and observed-to-canonical coverage between 0.3 and 1.2.

Repeated candidates were capped by PDB entry and anchor before coordinate processing to limit domination by highly repeated large assemblies. Assemblies with 3–10 and 11–24 protein chains formed the primary benchmark. Assemblies containing 25–60 chains were retained only for exploratory analysis, whereas assemblies containing more than 60 protein chains were excluded.

The complete filtering sequence reduced 91,859 Class A records to 70,830 after mapping and same-taxon requirements, 26,597 after PDB- and anchor-level caps, 11,504 after coordinate quality control, and a 6,051-record stratified mother pool. The primary 3–24-chain core contained 3,051 unique A–C noncontacts.

### Direct-partner selection and triplet formation

For each oriented A–C pair, candidate D proteins were required to have structural evidence of direct contact with A, defined using the same ≤8 Å minimum heavy-atom criterion. Contact evidence could originate from the biological assembly containing the A–C noncontact or from the aggregate PDB contact index. Candidate D proteins were prioritized for sequence resemblance to C using local BLOSUM62 alignment. Eligible triplets satisfied a retained similarity tier of at least 20% sequence identity with 50% bilateral coverage or a more stringent identity–coverage tier.

Among the 3,051 primary A–C pairs, 1,606 had at least one eligible D, yielding 3,365 A–D+/A–C− triplets. When multiple D candidates were available for one A–C pair, D-level ranking outcomes were averaged within that A–C pair before benchmark-level aggregation. The replicated structural subset required the A–C noncontact to be observed in at least two distinct PDB entries; 1,894 of the 3,051 primary A–C pairs met this criterion. Exact A–C direct-contact conflicts in the aggregate PDB pair index were removed.

We additionally performed a conservative provenance audit of D. A triplet was classified as verified same-assembly only when D could be linked to the biological assembly supporting the corresponding A–C noncontact and A–D satisfied the ≤8 Å contact criterion in that assembly. This identified 555 verified same-assembly triplets. The remaining 2,810 triplets used D candidates sourced from the aggregate PDB contact index. Because these latter candidates were not exhaustively reconstructed against every possible supporting assembly, this category does not establish that A–D contact occurred in a different assembly. The verified same-assembly subset was therefore used as a conservative sensitivity analysis rather than as an exhaustive partition of structural provenance.

### Sequence-model inference

We evaluated released checkpoints for PLM-Interact, D-SCRIPT, Topsy-Turvy and MINT, together with the classical SPRINT baseline^6–10^. PLM-Interact jointly tokenized paired canonical sequences, applied longest-first truncation to a maximum input length of 1,024 tokens and used the released ESM2/classifier state dictionary. D-SCRIPT and Topsy-Turvy used their released human checkpoints; pairs containing sequences beyond the supported 2,000-residue inference limit were excluded from model-specific analyses.

MINT used the released interaction encoder and pair-classification head. Because its official collator randomly crops chains exceeding 512 residues, every selected pair was evaluated in both sequence directions under five fixed crop seeds (13, 29, 47, 71 and 97). Reciprocal directions were averaged within seed, and the five seed-level scores were then averaged before triplet ranking.

SPRINT was evaluated using the official OpenMP implementation with default PAM120 parameters. Its positive-training set comprised the mapped D-SCRIPT human training interactions used for the corresponding classical baseline. Because SPRINT scores are unbounded, only within-model rankings and within-model score-distribution comparisons were interpreted.

For the complete structural benchmark, the primary endpoint was the probability that the direct partner outranked the assembly noncontact, \(P[s(A,D)>s(A,C)]\), with exact ties contributing 0.5. When an A–C pair had multiple eligible D candidates, the D-level ranking indicators were first averaged within that A–C pair and the resulting A–C-level values were then averaged across the benchmark. This macro-averaging prevented A–C pairs with more eligible D candidates from contributing disproportionate weight.

### Random-reference controls

Two distinct random controls were used and were not interchangeable.

The 2,000-pair random unlabeled reference set was sampled from the 5,968 A/C proteins represented in the 6,051-record structural mother pool using seed 20260806. Self-pairs and pairs present in the structural binding or noncontact indices were excluded. These pairs were not matched by species, sequence cluster or biological context and were used only as an unlabeled score-distribution reference.

The same-anchor random-control set was constructed separately from a 1,000-triplet structural pilot. For each anchor A, an unrelated partner R was selected from proteins represented in the pilot after excluding A, C and D and proteins belonging to the same MMseqs30 cluster as C or D. Selection used seed 20260814. The locked analysis contained 300 triplets from 266 unique anchors and was used only for the D>R, C>R and D>C comparison in Fig. 2.

### Representation geometry and PPI-adaptation analysis

We analysed frozen representations from ESM2, the released PLM-Interact-650M-humanV12 model and MINT^20^. For ESM2 and PLM-Interact, the two canonical sequences were supplied as the same concatenated pair input used by the released PLM-Interact pipeline (maximum length 1,024 tokens), and hidden_states[ℓ][:,0,:] provided the pair-CLS vector at layers 0–33. For MINT, each chain was cropped to at most 512 residues using fixed seed 13, residues excluding special tokens were mean-pooled separately for the two chains, and the two 1,280-dimensional vectors were concatenated. The matched MINT ESM2 control used the same cropped residues but processed the two chains independently without multimer attention. All 34 hidden-state layers (embedding layer 0 and transformer layers 1–33) were extracted before examining cluster-unseen performance. Layer 33 was the prespecified confirmatory readout; complete layer-wise profiles were exploratory. For same-anchor geometry, we calculated cosine similarities among A–D, A–C and A–R representations and summarized 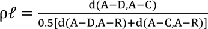, where d=1−cosine similarity.

### Cluster-disjoint representation probes

The 3,365 structural triplets were partitioned by MMseqs2 50%-identity families^41^. The full split contained 2,124 training-side triplets, 101 protein-unseen/cluster-seen triplets and 540 cluster-unseen triplets. An additional 277 mixed-cluster, 273 mixed-protein and 50 same-PDB triplets were excluded from the clean transfer evaluation. Iterative matched-random closure reduced the training-side pool from 2,124 to 1,865 train-eligible triplets.

Two frozen-representation probes were fitted. The context probe distinguished biologically plausible A–D/A–C relationships from matched A–R relationships. The contact probe distinguished A–D from A–C. Feature standardization parameters were estimated from the training set only. Labels were mapped to −1,+1, an intercept was added, and ridge regression used the prespecified regularization parameter λ=1; the same penalty was applied to the full coefficient vector. Probe outputs were treated as continuous scores rather than probabilities.

For ESM2, PLM-Interact and MINT, 1,865 training and 540 cluster-unseen triplets entered the common analysis. Cluster-unseen examples were not used to fit probe parameters or select the confirmatory layer. Exploratory layer-wise maxima were reported only in Supplementary analyses.

For the common AUROC probe analysis, uncertainty was estimated by 200 percentile bootstrap resamples over triplet IDs using seed 20260818; all scores belonging to the same triplet were resampled together. The point estimate was calculated on the complete evaluation set rather than as the bootstrap mean.

### Matched structural-negative fine-tuning

The same cluster split was used for the matched fine-tuning analysis. Structure-FT and Random-FT were constructed from the 1,865 train-eligible triplets and contained identical anchors, direct-contact positives, negative-protein multisets and row counts. Structure-FT retained the observed A–C structural pairing. Random-FT reassigned C proteins within taxon while excluding the original C, A, D and known structural contacts. Thus, the two arms differed only in negative-partner assignment.

Both arms initialized the released PLM-Interact checkpoint and used binary cross-entropy with AdamW, a backbone learning rate of 10−5, classifier-head learning-rate multiplier of 10, weight decay 0.01, batch size 8, gradient accumulation 4, 5% linear warm-up, maximum input length 1,024 and three epochs. Three seeds (20260816, 20260817 and 20260818) were trained, each completing 315 optimizer steps. Scores were averaged across seeds before final ranking.

A historically named 186-triplet internal_dev subset remained part of the fitting pool and was not used for hyperparameter selection, early stopping or model selection. It should therefore not be described as a held-out development set. Protein-unseen/cluster-seen and cluster - unseen evaluation definitions followed the cluster partition described above.

### ESMFold and AlphaFold 3 evaluation

A common 300-triplet multimodel subset comprising 256 unique anchors was used for direct comparison of the five sequence-based PPI predictors, ESMFold and AlphaFold 3. This subset is distinct from the 300-triplet/266-anchor same-anchor random-control set used in Fig. 2. All seven methods produced complete scores for all 300 triplets.

ESMFold used the facebook/esmfold_v1 checkpoint through EsmForProteinFolding^20^. Each dimer was represented as sequence A followed by a 25-glycine linker and sequence B, with four recycles and chunk size 64. The prespecified endpoint was negative mean inter-chain predicted aligned error after excluding linker positions, so larger values represented more confident inter-chain geometry. All 577 unique dimers completed, but chain pLDDT remained below 16 and the linker protocol did not form a reliable interface; it was retained as a PLM-driven folding control rather than interpreted as a validated complex model. AlphaFold 3 version 3.0.3 was run on isolated A–D and A–C dimers with one seed, five diffusion samples, Triton attention and maximum template date 30 September 2021^21^. The AlphaFold 3 endpoint was ipTM. The PDB evidence entries used in the 300-triplet comparison post-dated the template cutoff, but exact-pair historical exposure was not reconstructed; the analysis is therefore a benchmark comparison rather than a formal zero-shot test.

Confidence intervals for the seven-model common-subset comparison were obtained from 10,000 anchor-block bootstrap resamples over the 256 unique anchors.

### Negatome experimental-negative analysis

Negatome Manual and Manual-stringent records were mapped to canonical UniProt accessions and sequences^30,31,40^. The primary semantic analysis required two human canonical proteins, compatibility with the PLM-Interact input window and removal of exact undirected overlap with positive STRING v12 training pairs^42^.

The prospective assay ontology classified purified or reconstituted biophysical measurements, purified pull-down, far-western and specified in-vitro binding experiments as reconstituted experimental direct non-binding; Y2H, BRET, FRET, BiFC, PLA, crosslink and related cellular assays as cellular binary/proximity; and co-immunoprecipitation, AP–MS, TAP and affinity co-complex assays as co-complex association. The primary harmonized cohort contained 298 reconstituted-direct, 145 cellular binary/proximity and 119 co-complex pairs, with 22 unresolved methods retained only for audit.

The historical direct_binary definition comprised the 298 reconstituted-direct plus 145 cellular binary/proximity pairs (n=443) and was retained only as a sensitivity analysis rather than used for primary cross-model comparisons.

### Training-familiarity analysis and matched semantic comparison

The clean structural cohort required a 3–24-chain assembly, compatibility with the PLM-Interact input window, no exact STRING v12 positive-pair overlap and no targeted IntAct direct-positive conflict, yielding 2,480 A–C pairs^17^.

Each endpoint was searched with MMseqs2 against proteins present in positive training interactions using 30% query/target-style sequence coverage^41^. Endpoint familiarity was defined as the smaller of the best retained identities for A and C; endpoints without a retained hit were assigned zero for analysis and displayed separately where appropriate.

The adjusted logistic analysis used published PLM-Interact score ≥0.5 as the binary outcome and included endpoint familiarity, A–C sequence similarity, minimum heavy-atom distance, pair sequence length, assembly size, bridge-interface contacts, repeated-PDB support and human–human status. Continuous covariates were standardized. Confidence intervals used two-way cluster-robust covariance over the MMseqs50 families of A and C.

For the matched semantic comparison, experimental and structural pairs were restricted to human proteins, excluded exact STRING-positive overlaps and satisfied the PLM-Interact input window. Experimental reconstituted-direct pairs were matched without replacement to clean structural noncontacts on the two endpoint-to-training identities, log pair length and absolute log endpoint-length ratio. The largest prefix for which every post-match absolute standardized difference was ≤0.15 was retained. Uncertainty was estimated by paired bootstrap resampling.

### Manually reviewed ProtNegDB literature cohort

A separate validation cohort comprised 480 manually accepted negative PPI pairs from 362 PubMed/PMC publications drawn from the ProtNegDB literature stream. Reported protein or gene identifiers, assay methods and taxa were extracted and mapped within the reported taxon to canonical UniProt accessions. Exact mapped accession-pair overlaps with Negatome were removed before independent comparison.

Reliable taxon-aware mapping retained 271 pairs, 264 had locally available canonical sequences and 262 remained after removal of exact Negatome overlaps. The primary contemporary experimental direct non-binding cohort contained 104 pairs, comprising biophysical direct measurements and other direct-binding assays. Fifty-three of the 104 required no PLM-Interact sequence truncation.

Biophysical direct assays included SPR, ITC, BLI, MST and fluorescence anisotropy. Other direct-binding assays included appropriately configured pull-down, in-vitro binding and far-western experiments. Association/proximity assays were retained for secondary method-stratified analyses but were not included in the 104-pair primary direct cohort.

### Secondary direct-positive partner-ranking analysis

For each eligible reviewed negative A–C, we selected one same-taxon IntAct candidate D annotated with direct interaction type MI:0407. Selection first preferred candidates for which both A–C and A–D fit the 1,024-token input window, followed by candidates supported by more distinct positive publications and evidence rows. One deterministic candidate was retained per negative. The primary paired endpoint was P[s(A,D)>s(A,C)], with exact ties contributing 0.5. Wilson intervals summarized the main proportions, and publication-cluster bootstrap was used in sensitivity analysis. These D candidates were database-direct annotations and were not individually re-reviewed in the source publications.

### Cross-model negative-semantics comparison

To compare evidence semantics without imposing a shared numerical threshold across models, we calculated the probability that a randomly selected clean structural-noncontact score exceeded a score from an experimental direct non-binding pair, P(score(structural)>score(experimental)), with ties contributing 0.5.

The structural cohort contained 2,480 clean assembly noncontacts. The primary Negatome comparison used the harmonized 298-pair reconstituted-direct cohort. The ProtNegDB comparison used 104 experimental direct pairs for PLM-Interact, MINT and SPRINT. D-SCRIPT and Topsy-Turvy scored 98 of the 104 ProtNegDB pairs because six pairs contained sequences exceeding their released 2,000-residue inference limit.

Uncertainty for these probability-of-superiority estimates was obtained from 2,000 independent bootstrap resamples of the structural and experimental cohorts. The broader 443-pair historical Negatome direct/binary definition was analysed only as a sensitivity analysis.

### Statistical analysis

Exact score ties contributed 0.5 to all pairwise ranking probabilities. For the complete structural benchmark and structural sensitivity analyses in Supplementary Fig. S2a,d–f, D-level ranking indicators were first averaged within each unique A–C pair when multiple D candidates were available. Point estimates were then averaged across A–C pairs. Confidence intervals were calculated from 10,000 percentile-bootstrap resamples over A–C units.

For the score-threshold summaries in Supplementary Fig. S2c, fractions with PLM-Interact score ≥0.5 were calculated directly over pair rows, and 95% confidence intervals were obtained from 10,000 percentile-bootstrap resamples over pair rows. Other simple binomial proportions used Wilson 95% confidence intervals unless a panel-specific resampling procedure is stated.

The same-anchor random-control analysis used its locked 300-triplet/266-anchor set. Representation-probe AUROCs were calculated on the complete cluster-unseen set, with 95% percentile intervals obtained from 200 bootstrap resamples over triplet IDs; all observations belonging to the same triplet were resampled together.

Fine-tuning comparisons in Supplementary Fig. S5 used 10,000 anchor-cluster-block bootstrap resamples for aggregate transfer analyses; comparisons of change from the original model used paired resampling. The seven-method common-subset analysis used 10,000 anchor-block bootstrap resamples over the 256 unique anchors.

Training-familiarity regression used two-way MMseqs2 50%-family-clustered covariance over the A and C endpoint families. Familiarity-matched experimental–structural comparisons used paired bootstrap resampling. For cross-model comparisons of experimental direct non-binding and assembly noncontact, probability-of-superiority estimates were calculated with ties contributing 0.5 and uncertainty was obtained from 2,000 independent bootstrap resamples of the structural and experimental cohorts.

Primary proportions in the supportive direct-positive partner-ranking analysis used Wilson 95% confidence intervals; publication-cluster bootstrap was used as a sensitivity analysis. The cross-dataset Spearman correlation across the five PPI predictors was treated as a descriptive measure of model ordering rather than as a hypothesis test. All inferential tests were two-sided where applicable, and no significance threshold was used to select cohorts, model layers or reported panels.

## Data availability

The PDB, UniProt, STRING v12, IntAct and Negatome resources are publicly available from their providers. The ProtNegDB website is available at http://protnegdb.bjmu.edu.cn/. Neural checkpoints are available from the original PLM-Interact, D-SCRIPT, Topsy-Turvy and MINT releases; SPRINT source code is available from its original release.

## Code availability

Analysis, quality-control, model-inference and figure-rendering code will be released at https://github.com/chsmiss/protnegdb-benchmark.

## Acknowledgements

Not applicable

## Author contributions

D.Z. and H.C. conceptualized and designed the study. H.C., C.S. and conducted a data analysis. H.C. and C.S. wrote and edited the paper. C.Y. and C.W. gave important suggestions. All authors read and approved the final manuscript.

