## Supplementary Information for "Evidence-resolved benchmarks separate biological context recognition from direct-partner specificity in protein interaction models"

#### Affiliations

### Table of contents

#### Supplementary Figures


Supplementary Figure S1  Structural benchmark construction and quality control.

Supplementary Figure S2  Robustness of structural direct-partner benchmarking.

Supplementary Figure S3  Representation adaptation and same-anchor geometry.

Supplementary Figure S4  Sequence-cluster-disjoint split and transfer audit.

Supplementary Figure S5  Structural-noncontact fine-tuning and transfer robustness.

Supplementary Figure S6  AlphaFold 3 partner-ranking sensitivity.

Supplementary Figure S7  Negatome filtering and evidence-semantic sensitivity.

Supplementary Figure S8  Training-familiarity and explanatory-model robustness.

Supplementary Figure S9  Contemporary ProtNegDB validation and partner-ranking sensitivity.

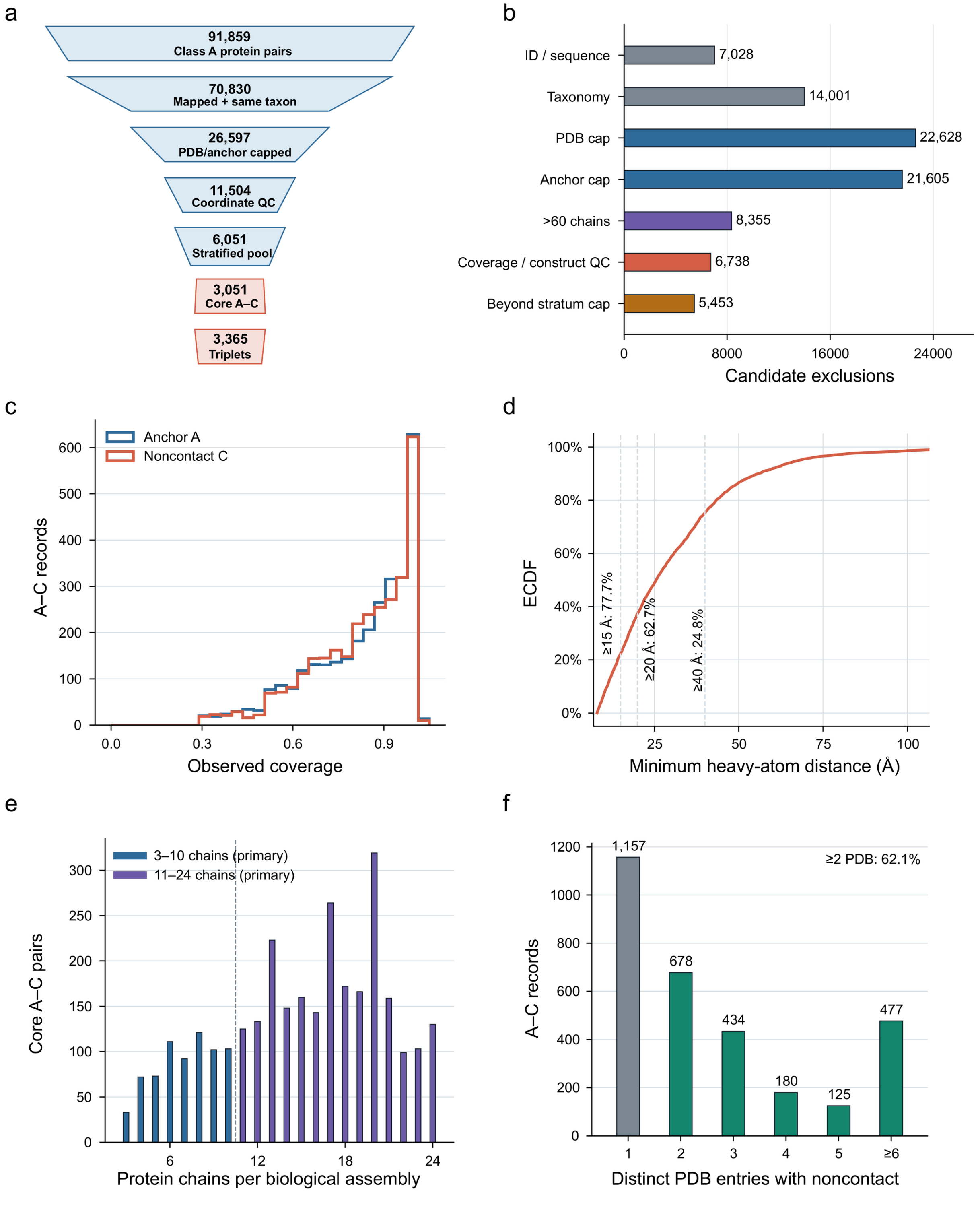


**Supplementary Figure S1 | Structural benchmark construction and quality control.** a, Filtering of the coordinate-resolved Class A branch to the primary A–C benchmark and same-anchor triplets. The final stage reports triplets rather than unique A–C pairs. b, Reasons for candidate exclusion across benchmark construction. Bars decompose the losses between successive stages of the filtering funnel in a. c, Observed-to-canonical sequence coverage of anchor A and noncontacting partner C in the 3,051 core A–C pairs, defined as the fraction of the canonical UniProt sequence represented by residues with resolved coordinates. d, Minimum heavy-atom distance between A and C in the supporting biological assembly. e, Assembly-size distribution of the primary benchmark, showing the prespecified 3–10- and 11–24-chain strata. Assemblies with more than 24 chains were retained only for exploratory analyses. f, Number of distinct PDB entries in which each core A–C pair was observed as a noncontact. In total, 1,894 of 3,051 pairs were supported by at least two PDB entries.

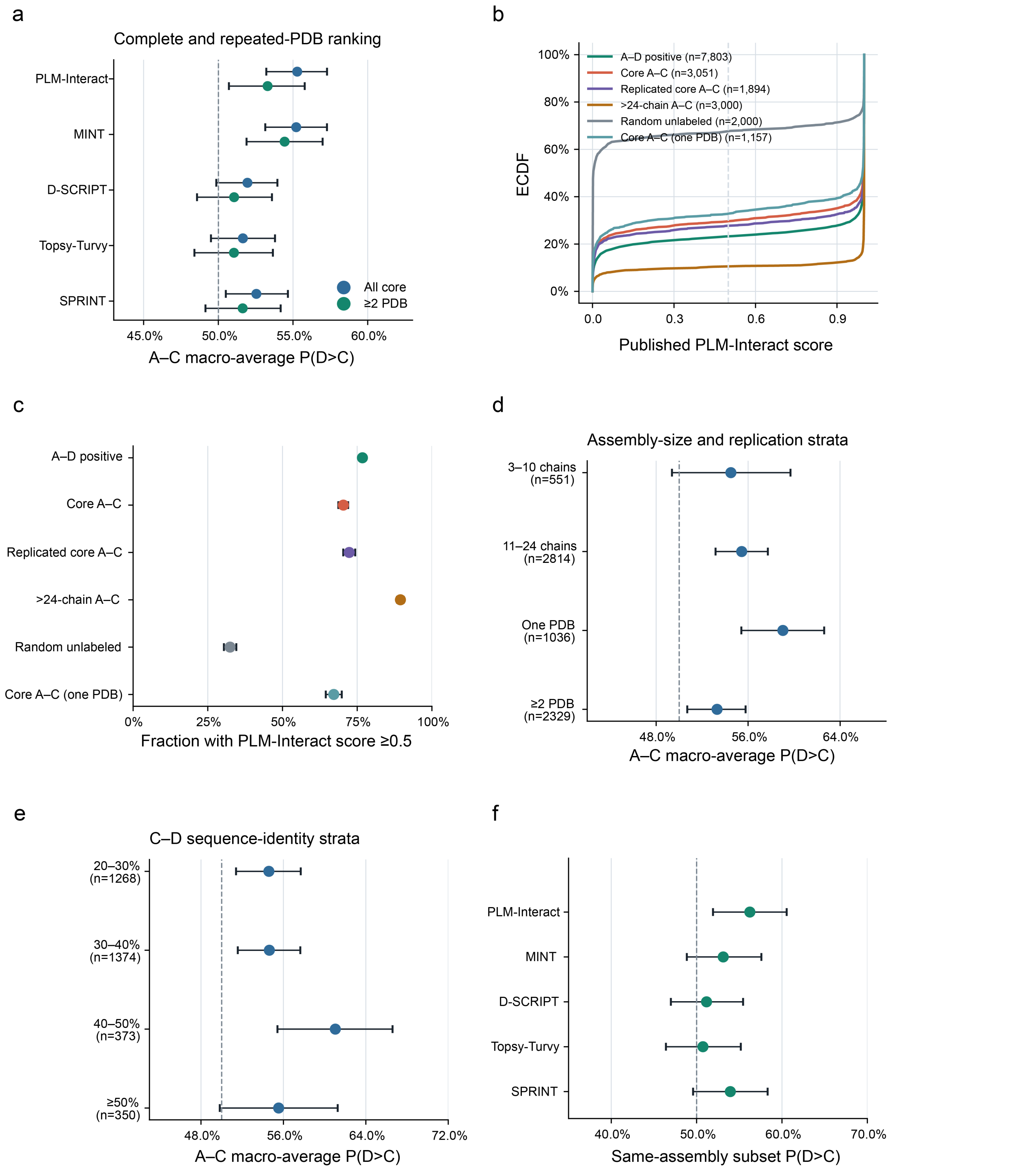


**Supplementary Figure S2 | Robustness of structural direct-partner benchmarking.** a, Direct-partner ranking in the complete and repeated-PDB subsets across five sequence-based PPI predictors. b, PLM-Interact score distributions for A–D direct-contact pairs, core A–C assembly noncontacts, repeated-PDB core noncontacts, exploratory >24-chain noncontacts, random unlabeled pairs and core noncontacts observed in a single PDB entry. c, Fraction of pairs with PLM-Interact score ≥0.5 in the same cohorts shown in b. The 0.5 cutoff is the published binary operating threshold of PLM-Interact and is used here only to summarize the prevalence of interaction-positive predictions. Repeated-PDB and one-PDB subsets stratify the core A–C noncontacts by structural recurrence, >24-chain A–C pairs are exploratory, and random unlabeled pairs provide a conventional reference. The cutoff is not used for direct-partner ranking. d, PLM-Interact direct-partner ranking across assembly-size and replication strata. e, Direct-partner ranking across C–D sequence-identity strata. f, Direct-partner ranking restricted to the 555 triplets in which A–D contact was verified in the same biological assembly supporting the A–C noncontact. The remaining 2,810 triplets used D candidates sourced from the aggregate PDB contact index; this designation does not establish that their A–D contacts occurred in a different assembly. The verified same-assembly subset is conservative rather than exhaustive. Ranking averages over D candidates within each A–C pair. Error bars are 95% percentile intervals from 10,000 bootstrap resamples over A–C units in a,d–f and pair rows in c. Model-specific sample sizes are provided in the source data; D-SCRIPT and Topsy-Turvy cover 547 of the 555 verified same-assembly triplets.

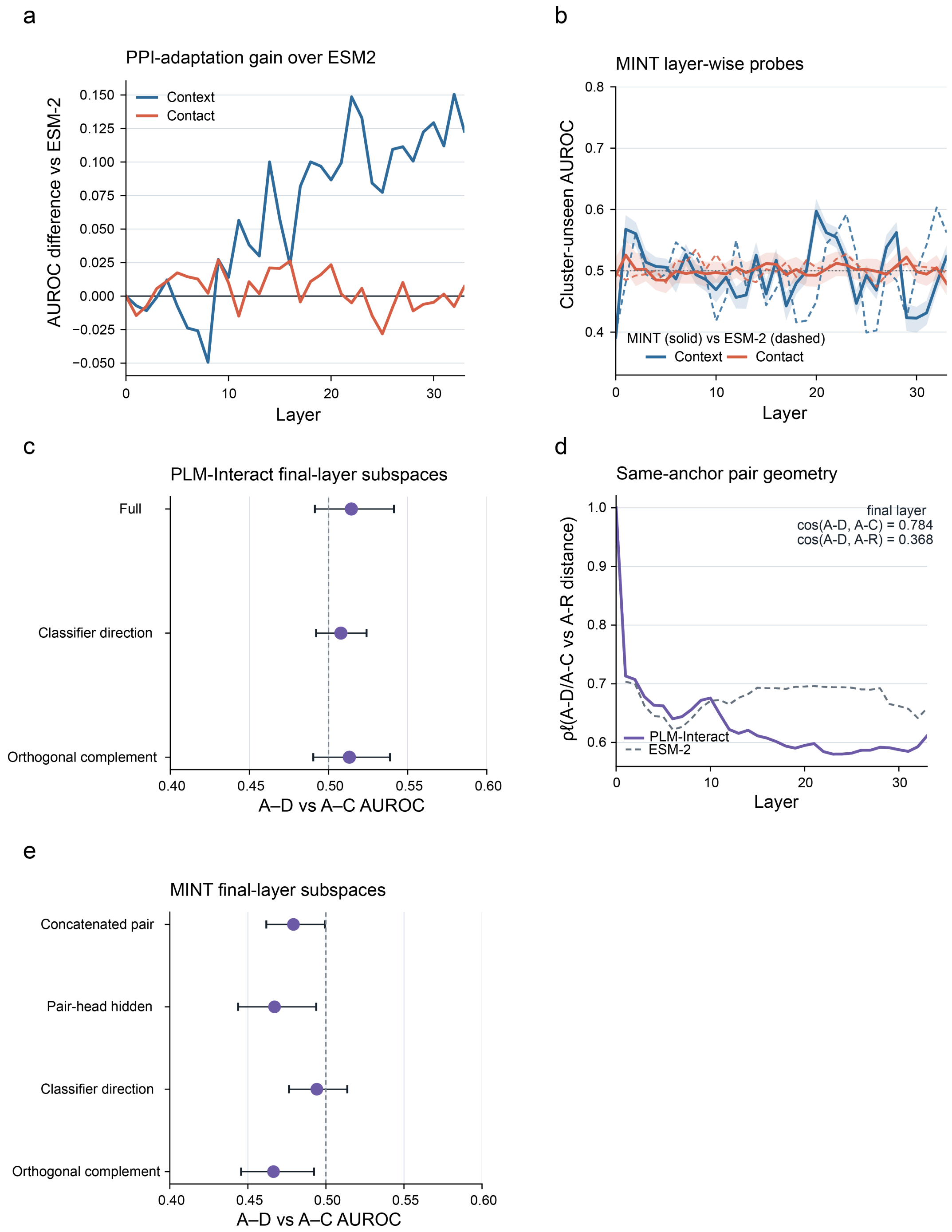


**Supplementary Figure S3 | Representation adaptation and same-anchor geometry.** a, Layer-wise difference in context and contact AUROC between PLM-Interact and ESM2. b, Layer-wise MINT probe performance and the matched ESM2 control. c, Final-layer PLM-Interact contact probes using the full representation, classifier direction and orthogonal complement. d, Same-anchor representation geometry summarized by $\rho\ell = d(A–D,A–C)/\{0.5[d(A–D,A–R)+d(A–C,A–R)]\}$, where $d = 1 - $ cosine similarity. e, Corresponding MINT final-layer subspace analysis. Probe analyses use 540 cluster-unseen triplets. Shading and error bars, where shown, are triplet-bootstrap 95% intervals. Layer-wise trajectories are exploratory, whereas the final-layer readouts are prespecified.

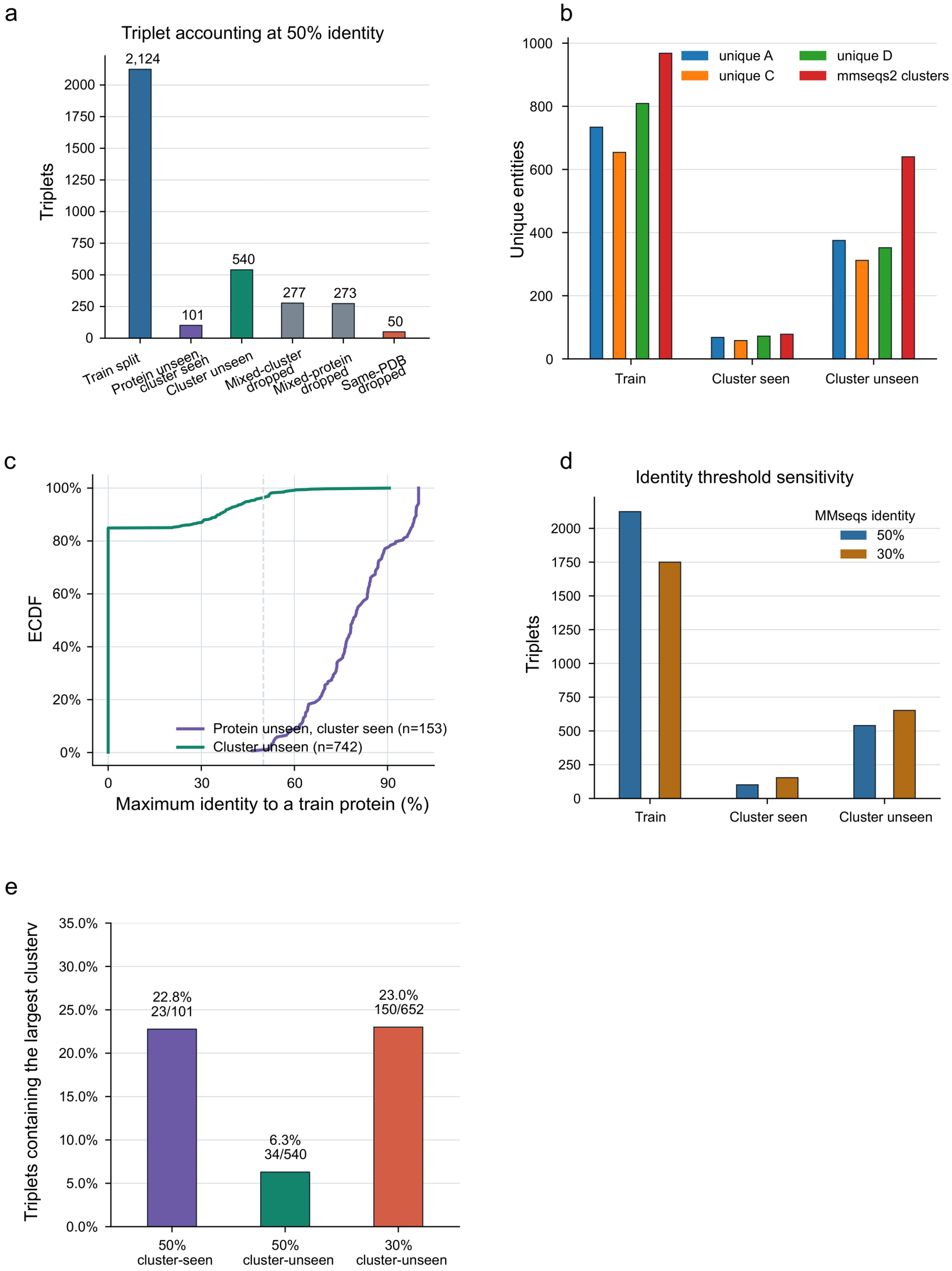


**Supplementary Figure S4 | Sequence-cluster-disjoint split and transfer audit.** a, Triplet accounting across training, protein-unseen/cluster-seen, cluster-unseen and excluded categories. b, Numbers of unique A, C and D proteins and MMseqs2 sequence clusters in the training, protein-unseen/cluster-seen and cluster-unseen subsets. c, Maximum retained sequence identity to a training protein for proteins in the transfer subsets; proteins without a retained hit are assigned zero. d, Split-composition sensitivity to MMseqs2 clustering thresholds of 50% and 30% sequence identity. e, Fraction of evaluation triplets containing the largest MMseqs2 cluster, counting each triplet once even when members of that cluster occur in multiple roles. Panel d evaluates split composition rather than retraining at both thresholds. The 2,124-triplet training-side pool precedes matched-random eligibility filtering to 1,865 train-eligible triplets.

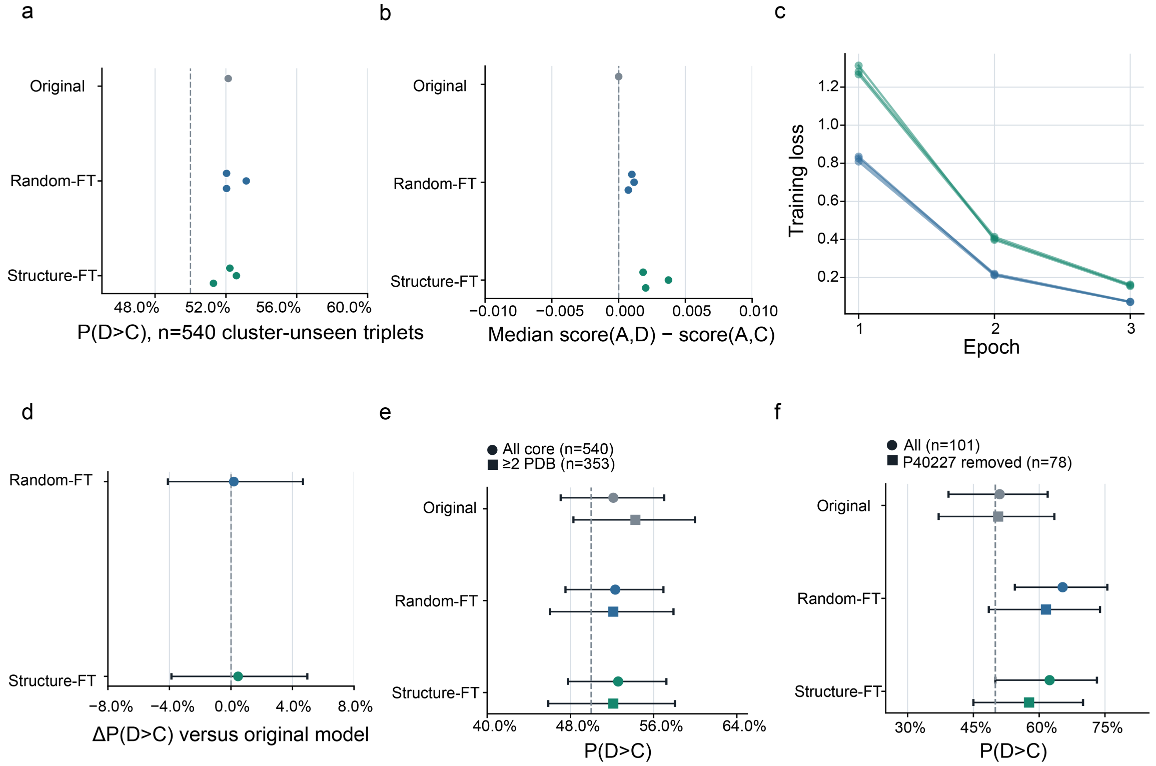


**Supplementary Figure S5 | Structural-noncontact fine-tuning and transfer robustness.** a, Per-seed cluster-unseen direct-partner ranking for the Original, Random-FT and Structure-FT models. b, Corresponding median A–D minus A–C score margins on a zero-centred axis; margins are descriptive and do not establish a meaningful method advantage. c, Training loss across the three fine-tuning seeds. d, Aggregate cluster-unseen ranking changes relative to the Original model. e, Complete and repeated-PDB comparisons within the cluster-unseen subset. f, Protein-unseen/cluster-seen performance before and after removal of the dominant P40227-containing MMseqs2 cluster. For aggregate comparisons, pair scores are averaged across seeds before ranking. Error bars in d-f use 10,000 anchor-cluster-block bootstrap resamples; d uses paired resampling. Structure-FT and Random-FT contain identical anchors, positive partners and negative-protein multisets and differ only in negative-partner assignment.

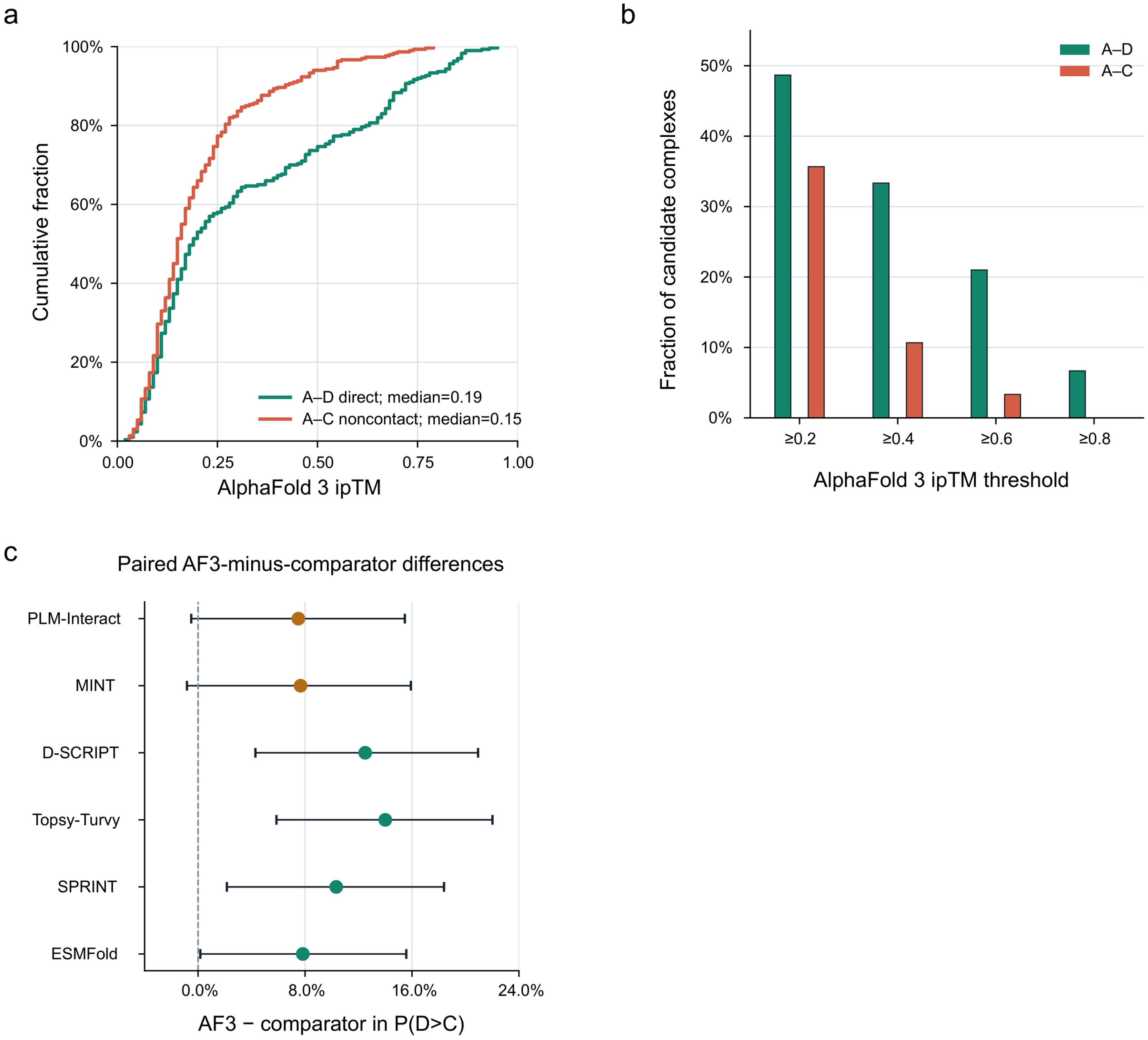


**Supplementary Figure S6 | AlphaFold 3 partner-ranking sensitivity.** a, AlphaFold 3 ipTM distributions for direct A–D and assembly-noncontact A–C candidate complexes. b, Fractions of A–D and A–C predictions exceeding the displayed ipTM thresholds. c, Paired differences in direct-partner ranking between AlphaFold 3 and each comparator. All panels use the common 300-triplet/256-anchor subset. Intervals in c are 95% anchor-block bootstrap confidence intervals. These comparisons do not establish formal zero-shot generalization.

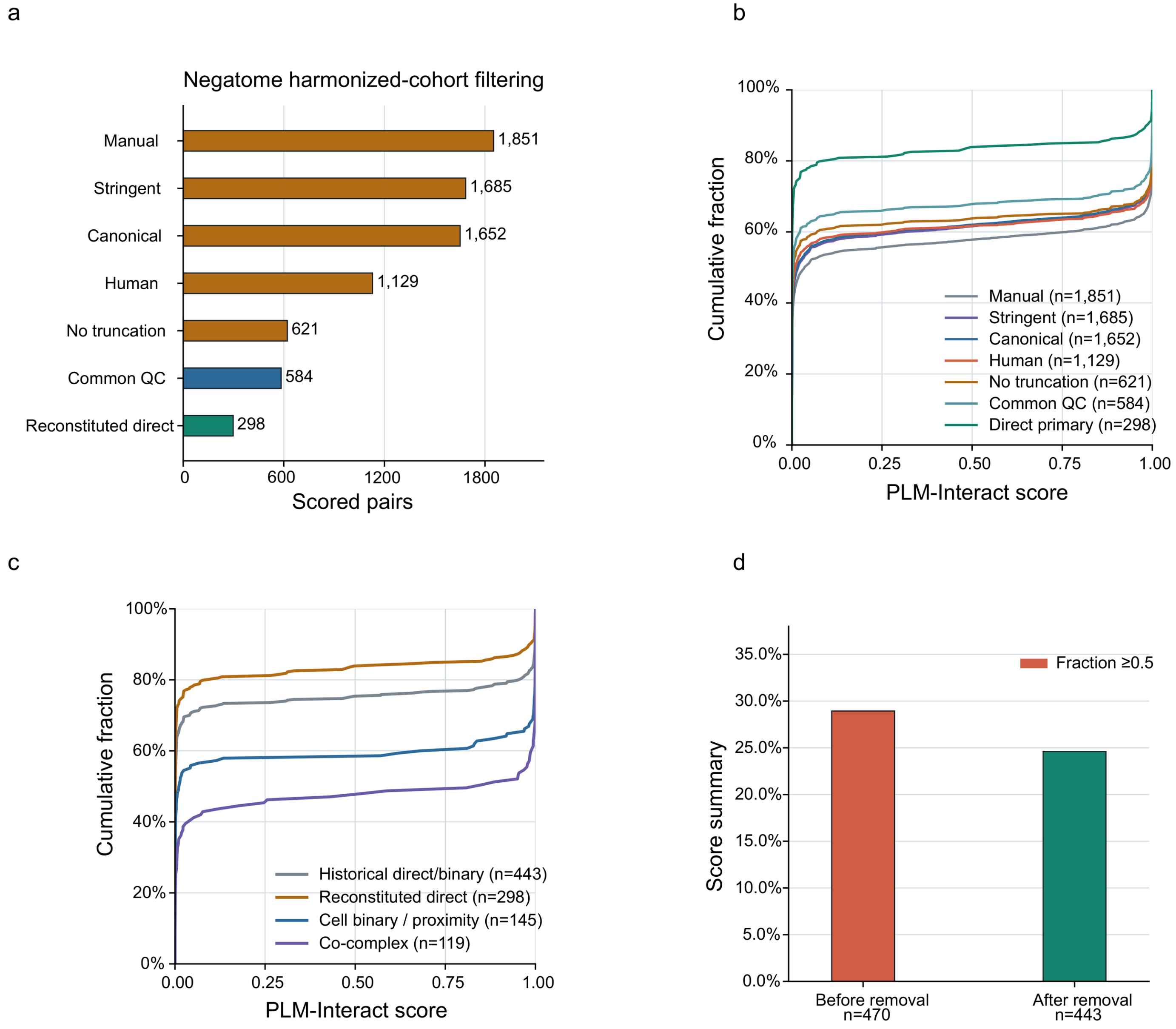


**Supplementary Figure S7 | Negatome filtering and evidence-semantic sensitivity.** a, Filtering of scored Negatome Manual records to common quality control and the harmonized reconstituted-direct cohort. b, PLM-Interact score ECDFs across successive quality-control stages. c, The historical direct/binary group compared with the harmonized reconstituted-direct, cellular binary/proximity and co-complex classes. d, Fraction of historical direct/binary pairs scoring ≥0.5 before and after removal of exact positive-training overlaps (470 and 443 pairs, respectively). The primary reconstituted-direct cohort contains 298 pairs; the historical 443-pair group additionally contains 145 cellular binary/proximity pairs. Changes across filtering stages reflect changes in cohort composition rather than causal effects of individual filters.

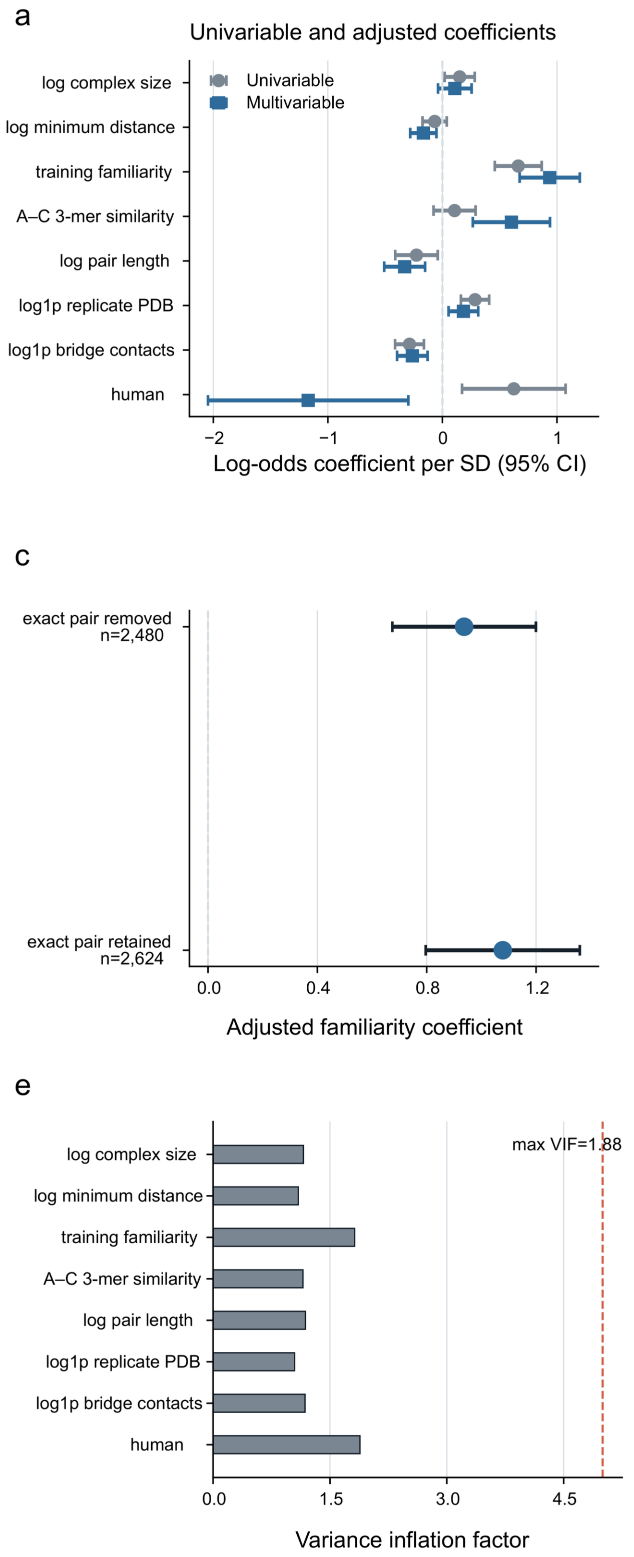


**Supplementary Figure S8 | Training-familiarity and explanatory-model robustness.** a, Univariable and multivariable associations with PLM-Interact score ≥0.5 among clean assembly noncontacts. b, Sensitivity to alternative endpoint-familiarity definitions. c, Sensitivity to handling of exact positive-training overlaps. d, Comparison of alternative uncertainty estimators. e, Variance-inflation factors for the adjusted model. Continuous predictors are standardized and human-human status is binary. Error bars show 95% intervals. These analyses assess associations rather than causal effects; the primary clean cohort contains 2,480 A–C pairs.

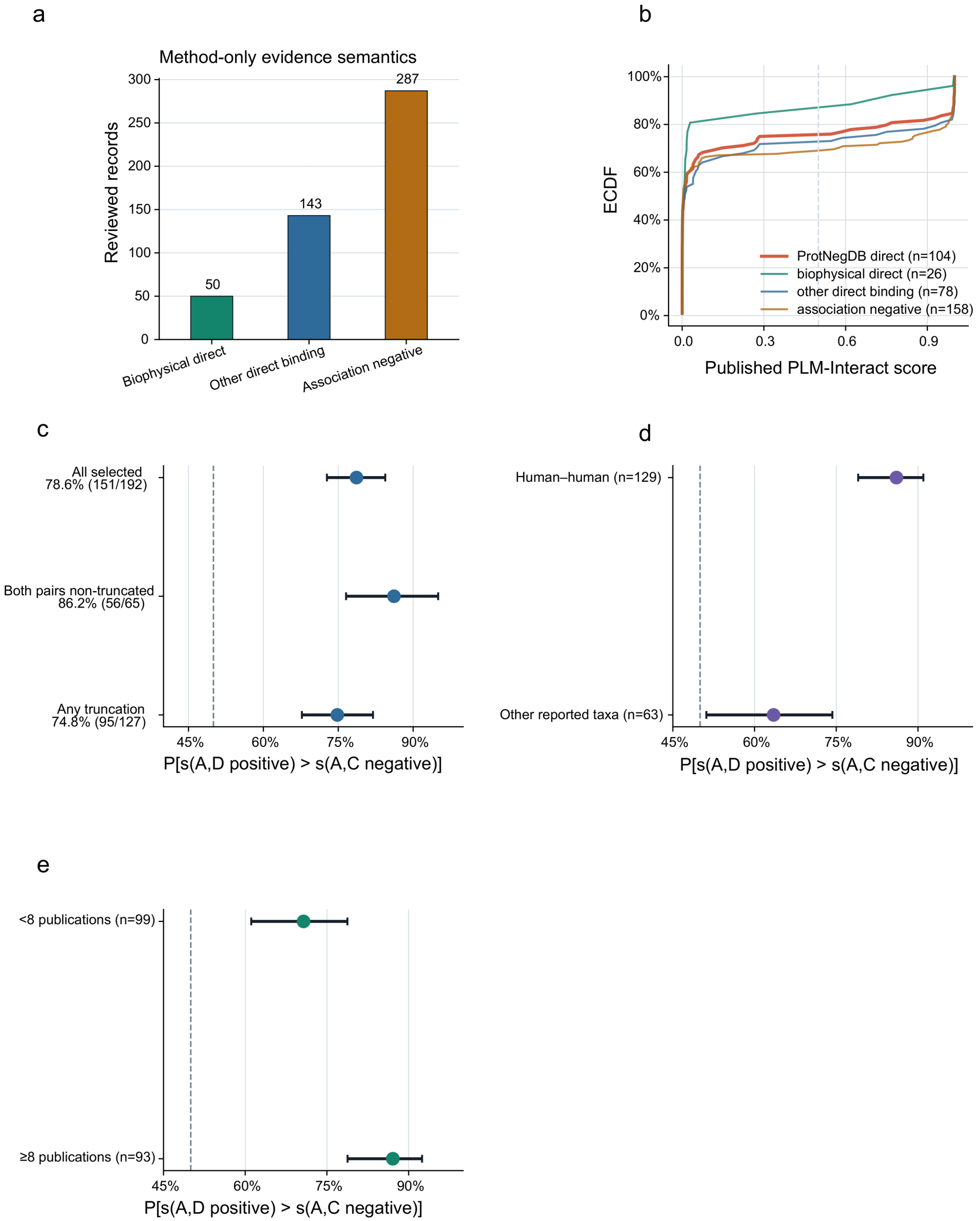


**Supplementary Figure S9 | Contemporary ProtNegDB validation and partner-ranking sensitivity.** a, Assay composition of the 480 manually reviewed negative pairs. b, Method-stratified PLM-Interact score distributions after mapping and removal of exact Negatome overlaps. c, Direct-positive partner-ranking sensitivity to sequence truncation; error bars are publication-cluster bootstrap 95% intervals. d, Partner-ranking results across species strata. e, Partner-ranking results stratified by publication support for the selected IntAct positive partner. Intervals in d and e are Wilson 95% intervals. The supportive ranking cohort contains 192 pairs; IntAct MI:0407 D candidates were not individually re-curated from their primary publications.
